# Combining Bacteriophage and Vancomycin is Efficacious Against MRSA biofilm-like Aggregates Formed in Synovial Fluid

**DOI:** 10.1101/2023.05.15.540793

**Authors:** Mariam Taha, Tia Arnaud, Tasia J. Lightly, Danielle Peters, Liyuan Wang, Wangxue Chen, Bradley W.M. Cook, Steven S. Theriault, Hesham Abdelbary

## Abstract

**Background:** Biofilm formation is a major clinical challenge contributing to treatment failure of periprosthetic joint infection (PJI). Lytic bacteriophages (phages) can target biofilm associated bacteria at localized sites of infection. The aim of this study is to investigate whether combination therapy of phage and vancomycin is capable of clearing S*taphylococcus aureus* biofilm-like aggregates formed in human synovial fluid.

**Methods:** In this study, *S. aureus* BP043, a PJI clinical isolate was utilized. This strain is a methicillin-resistant *S. aureus* (MRSA) biofilm-former. Phage Remus, known to infect *S. aureus*, was selected for the treatment protocol. BP043 was grown as aggregates in human synovial fluid. The characterization of *S. aureus* aggregates was assessed for structure and size using scanning electron microscopy (SEM) and flow cytometry, respectively. Moreover, the formed aggregates were subsequently treated *in vitro* with: a) phage Remus (∼10^8^ plaque-forming units (PFU)/mL), b) vancomycin (500 µg/mL), or c) phage Remus (∼10^8^ PFU/mL) followed by vancomycin (500 µg/mL), for 48 hours. Bacterial survival was quantified by enumeration (colony-forming units (CFU)/ mL). The efficacy of phage and vancomycin against BP043 aggregates was assessed *in vivo* as individual treatments and in combination. The *in vivo* model utilized *Galleria mellonella* larvae which were infected with BP043 aggregates pre-formed in synovial fluid.

**Results:** SEM images and flow cytometry data demonstrated the ability of human synovial fluid to promote formation of *S. aureus* aggregates. Treatment with Remus resulted in significant reduction in viable *S. aureus* residing within the synovial fluid aggregates compared to the aggregates that did not receive Remus (p < 0.0001). Remus was more efficient in eliminating viable bacteria within the aggregates compared to vancomycin (p < 0.0001). Combination treatment of Remus followed by vancomycin was more efficacious in reducing bacterial load compared to using either Remus or vancomycin alone (p = 0.0023, p < 0.0001, respectively). When tested *in vivo*, this combination treatment also resulted in the highest survival rate (37%) 96 hours post-treatment, compared to untreated larvae (3%; p < 0.0001).

**Conclusion:** We demonstrate that combining phage Remus and vancomycin led to synergistic interaction against MRSA biofilm-like aggregates *in vitro* and *in vivo*.

## Introduction

Prosthetic joint infection (PJI) is a devastating complication that can occur following total joint replacements (TJR). Current standard therapies for PJI have a failure rate reaching 30% (1). Treatment failure can lead to limb amputations and death (2). According to multiple national joint registries, the burden of PJI is a mounting crisis that will continue to impact healthcare systems (3). The current treatment protocol for PJI involves a shot-gun approach of administering long-term systemic antibiotics, in addition to multiple operations to debride the infected joint (3). However, systemic antibiotic therapies often fail due to the formation of biofilms. Biofilms are characterized as complex, interconnected communities that adhere onto surfaces within a self-produced matrix, offering protection from antibiotics and the host immune system (4-6).

*Staphylococcus aureus* is the most common biofilm-forming pathogen isolated from PJI patients with a prevalence of 30-40% of all PJI cultures (7). Enhancing the understanding of biofilm pathogenesis and development within synovial fluid, in-joint, can improve therapeutic approaches targeted against these biofilms. It is well documented that microbial biofilms adhered to implant surfaces play a key role in antibiotic tolerance, ultimately leading to treatment failure (8). Many recent studies demonstrate that *S. aureus* and other staphylococci are capable of organizing as floating biofilm-like aggregates in synovial fluid. This phenomenon has been reported *in vitro* and as well as in clinical synovial fluid samples (9-12). These biofilm-like aggregates elevate the minimum inhibitory concentration (MIC) of antibiotics necessary to control an infection and provide the bacteria with protection against clearance by the immune system (10, 13). Therefore, biofilm-like aggregates in synovial fluid are thought to play an important role in PJI treatment failure (10).

Failure to treat PJI-associated infections with conventional antibiotic therapy has motivated the search for alternative treatment strategies, such as bacteriophage (phage) therapy. Lytic phages are naturally-occurring viruses that specifically target bacteria. Phages can kill bacterial cells both on the surface and the interior of a biofilm (14, 15). Each phage has a specific affinity for infecting particular bacterial strains. This phage specificity is determined through recognition of bacterial surface proteins with their receptor-recognition and binding proteins.

Biofilm matrices possess water channel networks that allow nutrients, gases and water to reach bacterial cells through the different layers of the biofilm (16). It had been reported that some phages were capable of reaching the deeper layers of mature biofilms using these waternetworks to infect cells on the edge of the channels. This was proven using fluorescent-tagged phages. Biofilm architecture and density are contributing factors to the phage’s ability to diffuse efficiently through the biofilm (16). Moreover, some phages are capable of penetrating the biofilm and disrupting the biofilm matrix surrounding bacterial cells through the expression of enzymes such as depolymerases, lysins and DNases (17, 18).

Biofilm formation is a major clinical challenge contributing to treatment failure in patients afflicted with PJI. Lytic phages can target biofilm-associated bacteria at localized sites of infection. The aim of this study was to test the efficacy of phage therapy against biofilm-like aggregates of *S. aureus* formed in human-derived synovial fluid. Moreover, this study examined the ability of vancomycin, at a sub-lethal concentration, to synergistically interact with phage and boost the efficacy of the treatment *in vitro* and *in vivo*.

## Material and Methods

### Bacteria and phage

*S. aureus* BP043 (MRSA) was used in this work. It was previously acquired from The Ottawa Hospital Microbiology laboratory. The strain was collected from a PJI patient and was characterized as a biofilm-former (19). All bacterial cultures were incubated at 37°C, unless otherwise stated. *Silviavirus remus* or vB_SauM_Remus, formerly referred to as *Staphylococcus virus Remus,* is a member of the *Herelleviridae* family and will be referred to as: Remus, herein. It can be propagated on *S. aureus* PS47 in tryptic soy broth (TSB) (20). Remus was chosen for its efficiency of plating against *S. aureus* BP043 as it is equal to 1. This means that it lyses BP043 with similar efficiency as its host strain. Both phage Remus and *S. aureus* PS47 were obtained from Félix d’Hérelle Reference Center for Bacterial Viruses (www.phage.ulaval.ca). Tryptic soy agar (TSA) was used for underlays and overlays during phage titration.

### Phage propagation for *in vitro*

Propagation of Remus was carried out using *S. aureus* PS47 as the host bacterial strain in liquid media. An overnight culture of *S. aureus* PS47 was prepared by adding 2-3 colonies to 5 mL of TSB and incubated for 16-17 hours at 37°C, 200 rpm. A subculture was prepared by adding 100 µL of the overnight culture to 10 mL of TSB. The culture (10^7^ CFU/mL) was then, infected with 50 µL of Remus (at MOI =1) and incubated at 37°C at 200 rpm for ∼8 hours (until the culture is clear). Finally, the culture was filtered using a PES 0.45 µm syringe filter (Whatman Uniflo^TM^).

### Phage propagation and purification for *in vivo*

A late-exponential phase bacterial culture (10^8^ CFU/mL, OD_600_ = 0.8) of *S. aureus* PS47 was grown by inoculating 5 mL of TSB supplemented with 2 mM CaCl_2_ with a 1:100 dilution of an overnight culture of *S. aureus* PS47 and incubating at 37°C at 200 rpm. Then, 2.5 mL of the this culture was infected 1:1 (v/v) with Remus (∼10^2^ PFU/mL) at MOI =0.000001 and co-incubated at room temperature for 5 minutes to encourage phage adsorption. The phage-host mixture was then transferred to 100 mL of TSB with 2 mM CaCl_2_ and incubated at 37°C at 140 rpm, overnight. Phage lysate was processed by centrifugation at 4,000 x*g* at 4°C for 20 minutes and filtered using a 0.22 µm Nalgene™ Rapid-Flow™ bottle top filter (Thermo Fisher Scientific, Mississauga, Ontario, Canada). Phage was concentrated to 1 mL via ultrafiltration using Amicon™ 100 K Ultra-15 Centrifugal Filters (Sigma-Aldrich, St. Louis, Missouri, USA) at 4, 000 *xg* for 30 minutes. Purification of Remus by phase separation was carried out using Triton X-114, as previously described with modifications (21). The concentrated phage was treated with 1% (v/v) Triton X-114 and vortexed for 10 seconds. Samples were incubated on ice for 5 minutes, vortexed for 10 seconds, and incubated at 37°C for 5 minutes. Phase separation was carried out by centrifugation at 15,000 *xg* for 1 minute to collect the phage-containing, aqueous layer. This process was repeated 5 times. The aqueous layer was then washed, twice, at room temperature with 10 mL SM buffer via ultrafiltration, as described above, to remove residual Triton X-114. The purified phage sample was serially-diluted, 10-fold in SM buffer and titrated by spot plating on TSA in a soft agar overlay as previously described (22). Plaque forming units were determined after overnight incubation at 37°C.

### Synovial fluid

Human synovial fluid was either purchased from BioIVT, Westbury, NY, USA or collected from patients undergoing scheduled joint replacement surgeries at The Ottawa Hospital. Upon collection, the synovial fluid was centrifuged at 3,000 *xg* for 30 minutes to remove debris and cellular components. The supernatant was removed and filtered using a 70-µm strainer (Corning, USA). Sterility was assessed by plating 20 µL of the synovial fluid on TSA. The remaining synovial fluid was stored at −20°C.

### Biofilm-like aggregates formation

#### Bacterial inoculum

An overnight culture of *S. aureus* BP043 was prepared by inoculating 5 mL TSB with 2-3 colonies and incubating at 37°C with shaking (200 rpm) for 16-17 hours. Two mL aliquots of the overnight culture were centrifuged for 15 minutes at 3,000 *xg*. The supernatant was discarded and the pellet was resuspended in 2 mL of sterile, 1x PBS to give a final concentration of 1×10^9^ CFU/mL.

#### Synovial fluid

Bacterial aggregates were prepared in a 96-well plate in a 78% (v/v) synovial fluid solution comprised of: 140 µL synovial fluid, 20 µL of washed-bacterial cells and 20 µL of TSB. Synovial fluid was used at this high concentration (78%) to approximate the natural joint environment as closely as possible. The 96-well plate was then wrapped in parafilm and incubated, statically at 37°C for 24 hours to promote aggregate formation.

### Bright field and fluorescence microscopy

Pre-formed aggregates of BP043 in synovial fluid or planktonic cells in TSB from the overnight culture were collected (180 µL). Nucleic acid stain, SYTO9 Green (Invitrogen, ThermoFisher Scientific, Canada), at final concentration of 1 µg/mL was added to the cells, according to manufacturer’s instructions and were incubated for 30 minutes at room temperature. Cells were then washed 1x in HBSS (Hanks’ Buffered Saline Solution; Sigma, Oakville, Canada) then examined by adding 25 µL of the samples to the wells of a 12-well plate. Cover slips were added to the samples in the wells. Cells were imaged using a Cellomics, ArrayScan, VTI HCS Reader at 20x magnification.

### Flow cytometry

*S. aureus* BP043 grown as either planktonic cells (negative control) in TSB or pre-formed aggregates in synovial fluid were washed once in 1x PBS. Cells were fixed in 2% (v/v) paraformaldehyde for 10 minutes, then washed once in 1x PBS. Cells were stained with nucleic acid stain, SYTO9 Green according to manufacturer’s instructions. SYTO9 was added at final concentration of 1 µg/mL and samples were incubated for 30 minutes at room temperature. Cells were washed, once in 1x PBS, then examined using a BD LSRFortessa^TM^ flow cytometer with at least 10,000 events collected. A P2 gate was set to show SYTO9 green-positive bacterial aggregates with omission of unstained, single cells and debris.

### Scanning Electron Microscopy

The pre-formed aggregates were collected from 96-well plates. Aggregates were added to the fixative (4% paraformaldehyde and 2.5% glutaraldehyde) (v/v) and incubated for 1 hour at room temperature. Once the aggregates sedimented, the fixative was removed and samples were dehydrated in an ethanol gradient (20, 50, 70, 90 and 100%) followed by critical-point drying and microscopic examination.

### Biochemical Disruption Assay

Biochemical disruption assay was performed to investigate the extracellular matrix composition of *S. aureus* BP043 synovial-fluid aggregates and to quantitify the *S. aureus* cells contained within the aggregates. An overnight culture of BP043 was prepared in TSB and then washed, once in 1xPBS, as described earlier. *S. aureus* was added to pools of synovial fluid in a 12-well plate at a final volume of 2 mL/well. Synovial fluids were at a final concentration of ∼78%. Aggregates were encouraged to form at 37°C for 24 hours, statically. Proteinase K (ThermoFisher, Canada), at a final concentration of 150 µg/mL, or 1x PBS (control) were added to the preformed, synovial fluid aggregates. After 1 hour of incubation at 37°C, the aggregates were vortexed, serially-diluted and plated on TSA plates for enumeration (23). DNase I (Sigma, Oakville, Canada) was also used against *S. aureus* BP043 synovial fluid aggregates at a final concentration of 0.5 mg/mL (dissolved in 2 mM MgCl_2_). The Dnase I-and 1x PBS (control)- treated aggregates were incubated for 24 hours at 37°C (23, 24). Viable bacterial counts were enumerated as described above. Three to four biological replicates (3-4 pools of synovial fluid) were used with 2-3 technical replicates for each.

### Fluorescent Microscopy

Three fluorescent stains were used for the identification of the biofilm matrix composition. In particular, SYTO9 green-fluorescent was used to detect cellular biomass detection. Wheat germ agglutinin (WGA) Alexa Fluor® 350 conjugate detects carbohydrates, specifically bindingN-acetylglucosamine (GlcNAc) and N-acetylneuraminic acid residues(25). GlcNAc is a main component of poly-*β*-(1,6) N-acetyl-D-glucosamine (PNAG), hyaluronic acid, bacterial cell wall peptidoglycan and teichoic acid(26-29). FilmTracer™ SYPRO® Ruby, stains most classes of proteins(30). All the dyes were obtained from (ThermoFisher Scientific, Canada) and were used according to the manufacture’s recommendations. Briefly, the pre-formed aggregates were collected from the 96-well plate and washed, once with HBSS. SYTO9 was added and incubated for 15 minutes at a final concentration of 1 µg/mL. The aggregates were then removed and washed, once in HBSS. SYPRO Ruby was added at 1:1 ratio. After 30 minutes of incubation, the aggregates were washed, once with HBSS, and WGA was added at final concentration of 5 µg/mL. The aggregates were washed, twice in HBSS, added to a glass slide and left to air-dry, prior to examination using Zeiss AxioObserver 7 with Apotome2 microscope (20x objective). Each incubation period was at room temperature in the dark. Prolong^TM^ Gold Antifade Mountant (ThermoFisher Scientific, Canada) was used.

### Phage Remus lytic activities

The lytic activities of Remus against planktonic cultures of BP043 were assessed at different titers. A late-exponential phase bacterial culture (∼ 1 x 10^8^ CFU/mL) of BP043 was prepared by inoculating 5 mL of TSB with a 1:100 dilution from an overnight culture, then incubation at 37°C with 200 rpm shaking. A 96-well microtiter plate, containing 160 µL of TSB per well, was then inoculated with 20 µL of bacterial cells and 20 µL of diluted Remus (10^9^ – 10^2^ PFU/mL), corresponding to MOIs of: 10, 1, 10^-1^, 10^-2^, 10^-3^, 10^-4^, 10^-5^, 10^-6^. A bacterial control was prepared by inoculating a column of wells with 20 µL of SM buffer in place of phage. A media control was prepared by inoculating a column of wells with 180 µL with TSB and 20 µL of SM buffer. The optical density was then measured at 600 nm every 120 minutes at 37℃ for 48 hours. Two biological replicates were performed with six technical replicates for each MOI.

### Vancomycin

Vancomycin was chosen, since it is the first-line antibiotic treatment in orthopedic surgery against MRSA (31, 32). Vancomycin (Sigma-Aldrich, Oakville, Canada) was dissolved in sterile water and diluted to a working concentration of 500 µg/mL. The vancomycin concentration of 500 µg/mL was selected to conduct the experiments, representing 250x the MIC (MIC = 2 µg/mL) a sub-Minimum Biofilm Eradication Concentration value for strain *S. aureus* BP043 (MBEC = 2500 µg/mL) (19).

### *In vitro* Phage Remus against treatment of aggregates

The lytic activities of Remus were assessed against pre-formed aggregates of *S. aureus* BP043 in synovial fluid. In 96-well plates, 100 µL of Remus was added to pre-formed aggregates of *S. aureus* BP043 (∼ 4×10^8^ CFU/mL or 8×10^7^ CFU) at different concentrations: 10^9^ - 10^6^ PFU/mL. TSB was added (100 µL) to the pre-formed aggregates, as a control (no treatment). The plate was incubated for 48 hours under static conditions, at 37°C. Then, proteinase K (ThermoFisher, Canada), at a final concentration of 150 µg/mL, was added to the aggregates and incubated 1 hour at 37°C. The bacterial aggregates were vortexed, vigorously, sonicated in a water bath for 10 minutes, then serially-diluted and plated on TSA plates to assess bacterial survival in CFU/mL.

### *In vitro* Phage Remus and vancomycin treatment of aggregates

The pre-formed aggregates were exposed to: a) Remus (∼ 10^8^ PFU/mL) for 48 hours, b) vancomycin, 500 µg/mL for 48 hours, c) phage Remus for 24 hours, followed by vancomycin for 24 hours, or d) no treatment (control) for 48 hours under static conditions, at 37°C. Then, aggregates were processed as described above, assessing bacterial survival in CFU/mL.

### Phage Remus and vancomycin interaction

To determine the type of the interaction between Remus and 500 µg/mL vancomycin against the preformed biofilm-like aggregates, the coefficient of drug interaction (CDI) was calculated using bacterial counts (CFU/mL), according to the following equation(33, 34):

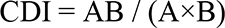

Where,

A, ratio of bacterial counts of phage treatment to bacterial counts of control group;

B, ratio of bacterial counts of vancomycin treatment to bacterial counts of control group;

AB, ratio of bacterial counts of combination treatment (phage and vancomycin) to bacterial counts of control group

If CDI is:

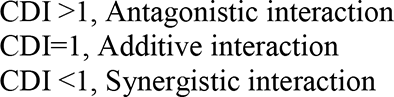

CDI < 0.7 indicates significant synergistic interaction

### Phage Remus viability and replication in synovial fluid

The capacity of Remus to survive and propagate within synovial fluid was analyzed. *S. aureus* BP043 aggregates were formed in synovial fluid and infected with Remus, as described above. After 48 hours, samples were serially-diluted in 1x PBS and plated onto lawns of *S. aureus* PS47 in TSA, as previously described (35). Plates were incubated, overnight at 37°C and plaques were counted. The initial concentration of Remus was 1×10^8^ PFU/mL. This was repeated 4 times with two technical replicates, each.

### Phage-resistance

The rise of phage-resistance sub-populations was monitored by evaluating the efficiency of plating (EOP) as previously described (36). Briefly, *S. aureus* aggregates were grown in synovial fluid, treated with Remus and then plated for viable bacterial counts, as described above. We selected four random *S. aureus* BP043 isolates that grew on TSA, post-Remus infection or no Remus treatment (control), then streaked onto TSA plates. After an overnight incubation at 37°C, a second subculture was performed by picking isolated colonies from the plates and re-streaking onto new TSA plates followed by overnight incubation. Liquid cultures were then prepared by adding single colonies from the second subculture to 5 mL of TSB and incubated at 37°C with 200 rpm shaking. The overnight, liquid culture was used to create a lawn of bacteria on TSA for each isolate and 10 μL of the serially-diluted Remus lysate was spotted. EOP was calculated by dividing Remus titer on the tested *S. aureus* BP043 by Remus titer on the ancestral *S. aureus* BP043. Two synovial fluids were used and four *S. aureus* isolates were checked for resistance per synovial fluid.

### *In vivo Galleria mellonella* bacterial infection model and phage treatment

*Galleria mellonella* larvae were used to assess the *in vivo* application of Remus and/or vancomycin to treat *S. aureus* infections with either: planktonic cells or synovial fluid-induced aggregates. ***Galleria mellonella Larvae****. G. mellonella* larvae were obtained from Serum Therapeutics Inc. (Edmonton, Alberta, Canada). Upon delivery, larvae were maintained in the dark, on-feed at 24°C. At least one day prior to use, larvae weighing ≥ 170 mg were removed from feed and starved at 24°C. Groups of ten larvae were selected based on similar weight and, lacking signs of melanization and pupation. ***Establishing infection.*** An overnight culture of BP043 was prepared and diluted to OD_600_ = 0.75 in TSB. The cells were then harvested by centrifugation at 4,000 *xg* for 10 minutes at 4°C and resuspended in either: a) 500 µL of Ringer’s solution for planktonic cells (control) or, b) 500 µL of 10% (v/v) human synovial fluid (BioIVT, Westbury, NY, USA) prepared in sterile Ringer’s solution (37). Then, cells were left to incubate, statically at room temperature (23°C) for 1 hour to promote *S. aureus* aggregation (37). Larvae were then injected in the left-hind proleg with 10 µL of ∼ 10^9^ CFU/mL (∼ 10^7^ CFU/inoculation) of aggregated or planktonic *S. aureus* cells and incubated for 1 hour to establish infection at room temperature. The injected-bacterial concentration used to initiate an infection was verified by serial dilutions and plating onto TSA for CFUs determination after overnight incubation at 37°C. The *S. aureus* inoculum delivered per larvae was around 2 x 10^7^ CFU for planktonic cells and 1.7 x 10^7^ CFU for aggregates. ***Treatment***. After incubation, single or consecutive treatments of Remus phage (10^8^ PFU, one dose) and/or vancomycin (5 µg, one dose) were delivered in the right-hind leg and second-to-last left-hind leg, respectively. Consecutive treatments involved administering Remus first, followed by vancomycin. SM buffer was administered as phage-treatment control and sterile distilled water (ddH_2_O) was used as vancomycin-treatment control. Treatment groups of larvae received one of the following treatments, a) control synovial fluid: SM+ ddH_2_O, b) control: PBS + Remus + vancomycin, c) control *S. aureus*: SM+ ddH_2_O, d) *S. aureus*: Remus + ddH_2_O, e) *S. aureus*: Remus + vancomycin, f) *S. aureus*: SM + vancomycin. Larvae were incubated at 37°C and monitored for mortality by daily scoring for activity for 96 hours, post-treatment. Three biological replicates were performed with ten larvae per group. Larvae were scored for death, based on lack of response to stimuli (score of 0) according to the *G. mellonella* Health Index Scoring System (Table S1)(38). Larvae showing signs of pupation were censored from the data.

### Ethical Approval

Ethical approval was obtained from Ottawa Health Science Network Research Ethics Board to collect both; bacterial strains obtained from PJI patients and synovial fluid collected from patients going through scheduled joint replacement surgeries.

### Bioinformatic analysis

Phage Remus was previously sequenced (20). We examined its genome for putative tail fibers encoded in the virion morphogenesis module. The coding domain sequences (CDS) were translated and analyzed using InterProScan plugin for Geneious 2021.2.2 to identify functional domains (39, 40). InterProScan ran using the following applications: CDD, Coils, Gene3d, HAMAP, MobiDB-Lite, Panther, PfamA, Phobius, PIRSF, PRINTS, PrositePatterns, PrositeProfiles, SFLD, SignalP, SignalP_EUK, SignalP_GRAM_NEGATIVE, SMART, SuperFamily, TIGRFAM, TMHMM (39, 40). Further investigation into proteins of interest was completed using PHYRE2 to model the proteins against protein data bank (PDB) templates.

### Statistics

Bacterial counts (CFU/mL) were converted to log_10_ scale. To compare the average size of aggregates (median fluorescence intensity of forward scatter) in TSB in comparison to synovial fluid: an unpaired, two-tailed t-test was utilized. The same test was used to compare the prevalence of aggregates formation expressed as percentages in TSB and compared to synovial fluid. To compare bacterial load after exposing aggregates to 1x PBS (control), and compare it to treatment of proteinase K or DNase I: an unpaired, two-tailed t-test was utilized. One-way ANOVA and post-hoc Tukey’s multiple comparisons test were performed to evaluate the efficiency of phage and vancomycin treatment regimes in reducing viable bacteria in biofilm-like aggregates in comparison to the control. One-way ANOVA and post-hoc Tukey’s multiple comparisons test was also used to compare the OD_600_ of planktonic cultures exposed to different phage concentrations at 48 hours. *P*-value<0.05 was considered to be significant. Kaplan-Meier survival curves and log-rank tests were used to evaluate the survival rates and statistical significances, respectively, of *in vivo* phage and vancomycin treatment in *G. mellonella*. *P*-value <0.001 was considered statistically significant between treatment groups. Analyses were performed using GraphPad Prism (GraphPad^TM^ Software, USA).

## Results

### Characterizing *S. aureus* aggregates formation and structure in human synovial fluid

The ability of human synovial fluid (∼78%) to influence *S. aureus* BP043 (MRSA PJI isolate) to form biofilm-like, aggregates compared to TSB media was explored. The presence of aggregates was examined macroscopically and microscopically (fluorescent microscopy and SEM). Aggregate size and abundance were assessed using flow cytometry. As shown in Figures 1 and 2, human synovial fluid at ∼78%, indeed supported aggregate-formation compared to TSB. The aggregates were visible macroscopically to the naked-eye (Figure 1, A). The aggregates that formed in synovial fluid dissipated after treatment with phage Remus (Figure 1, A). Images of bright field and fluorescence staining with SYTO9 (Figure 1, B) confirmed aggregates formation of BP043 in synovial fluid versus single, planktonic cells grown in TSB (negative control). SEM image of the aggregates displayed *S. aureus* clumping and filament-like structures, connecting the bacterial cells (Figure 1, C). Flow cytometry evaluation of the aggregates size, indicated the presence of larger *S. aureus* aggregates in synovial fluid than in TSB (Figure 2, A and C). Median fluorescence intensity of forward scatter (MFI-FSC) was significantly higher in synovial fluid compared to TSB, (p = 0.04). Moreover, figure 2, B presented more than 3x higher percentage of abundance of aggregates in synovial fluid (21%) versus that of TSB (5.6%, p = 0.02).

**Figure 1.**
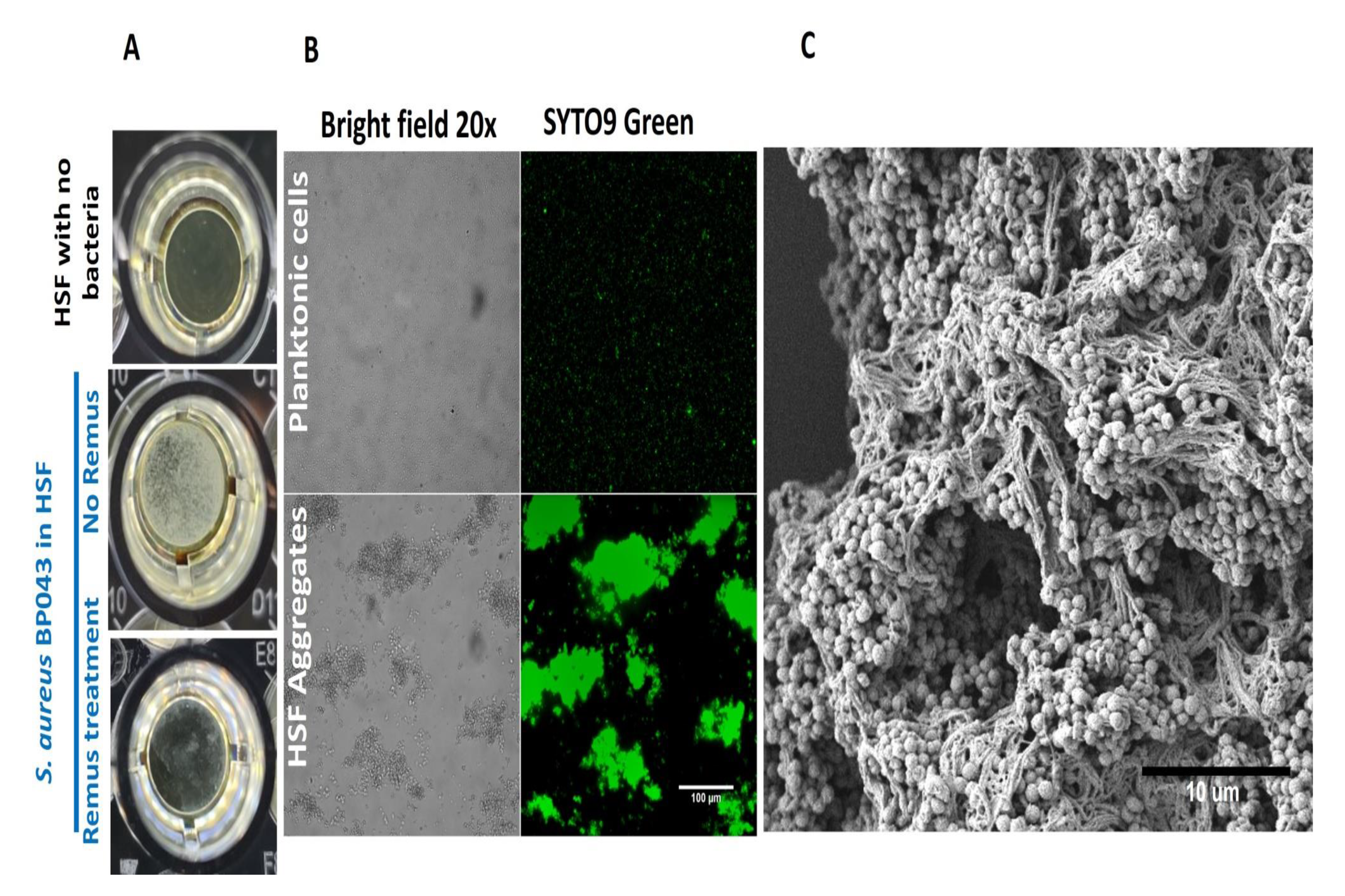
*S. aureus* BP043 aggregate formation in human synovial fluid (HSF) after incubation for 24 hours at 37°C. **A)** Macroscopic images of the *S. aureus* aggregates forming in human synovial fluid in 96-well plate, with and without Remus treatment. **B)** Images taken with bright field and fluorescence microscopy (SYTO9 nucleic acid stain) at 20x for *S. aureus* planktonic cultures grown in TSB or *S. aureus* grown as aggregates in human synovial fluid (HSF). **C)** Scanning electron microscopy image of *S. aureus* human synovial fluid aggregates.

**Figure 2.**
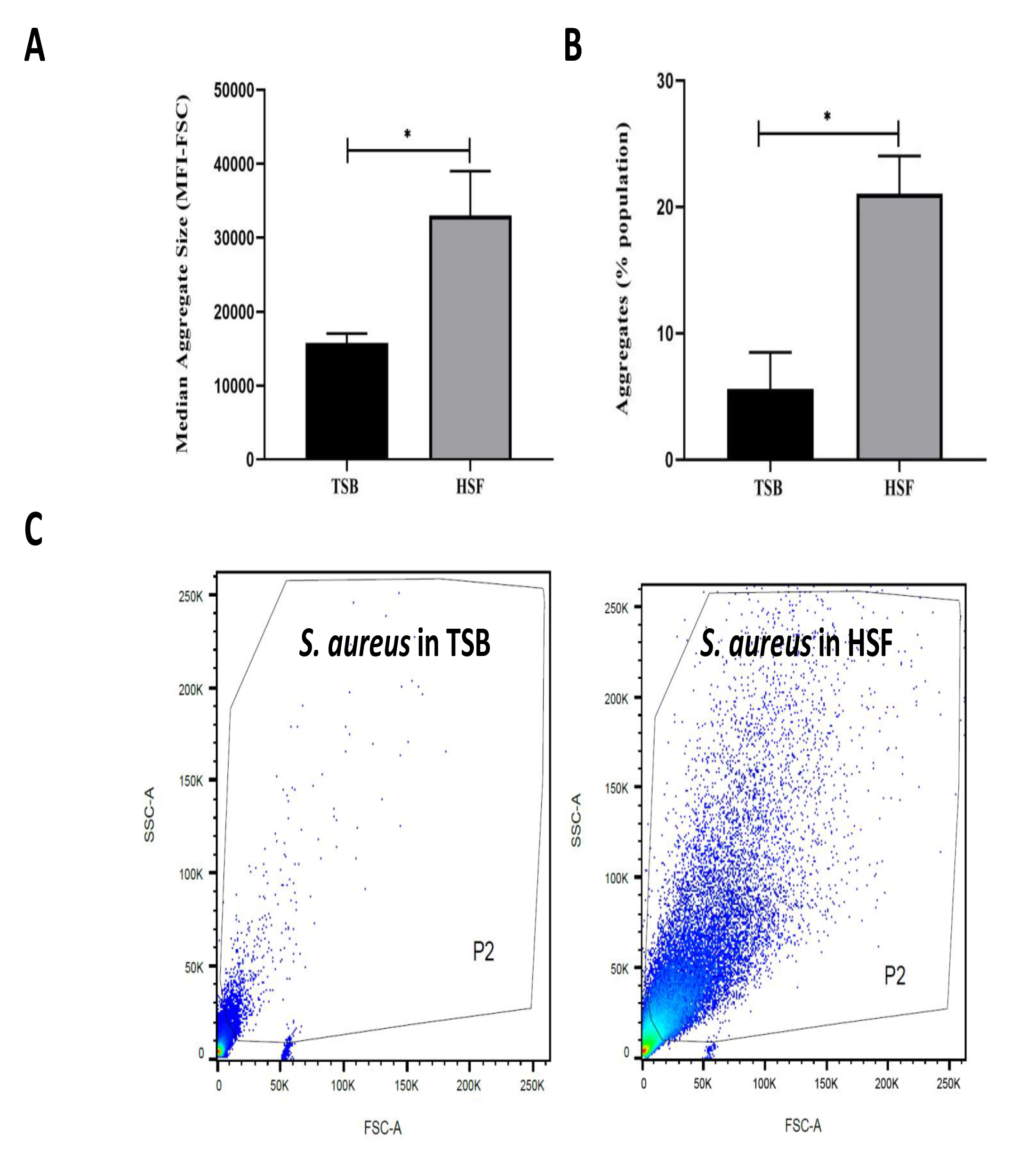
Human synovial fluid (HSF) enhances aggregate formation compared to TSB. Flow cytometry data shows **A)** bigger (median fluorescence intensity of forward scatter, MFI-FSC) and, **B)** more aggregates formation in HSF compared to TSB. **C)** P2 gate was set to show SYTO9 green positive bacterial cells. N = 3, mean ± SE. Unpaired, two-tailed t-test was utilized to determine statistical difference, *p < 0.05.

### *S. aureus* synovial fluid aggregates composition

To determine the composition of the human synovial fluid-induced aggregates, biochemical disruption assays using proteinase K and DNase I were conducted and bacterial viability was assessed. Moreover, the aggregates composition was determined by staining for polysaccharides (WGA) and proteins (SYPRO Ruby) by fluorescent microscopy. The addition of proteinase K led to the destruction of aggregates which was visually-observed (Figure 3, A). Proteinase K treatment also released bacterial cells from the aggregates which was defined by the increase in the CFU (increased by 1.3 log unites), compared to the untreated aggregates (p = 0.017, Figure 3, C). This observation suggests that the aggregates are rich in protein. In contrast, there was no macroscopic destruction of synovial fluid-induced aggregates after adding DNase I (Figure 3 B). Also, the DNase I treatment did not affect the overall bacterial load, indicating that the extracellular DNA has a minimal contribution to the aggregate stability (p = 0.85, Figure 3 B and D). Fluorescence microscopy images (Figure 4) of SYTO9-stained cells, displayed bacterial cellular biomass. A strong signal and large area of coverage with SYPRO Ruby (red color) also indicated the abundance of proteins contained within the aggregates. WGA-stain (blue color) was also noted but at a lower intensity, suggesting a lesser involvement of N-acetylglucosamine in aggregate composition. This pattern aligns with previously published data using SYPRO and WGA for synovial fluid aggregates (10, 13, 37).

**Figure 3.**
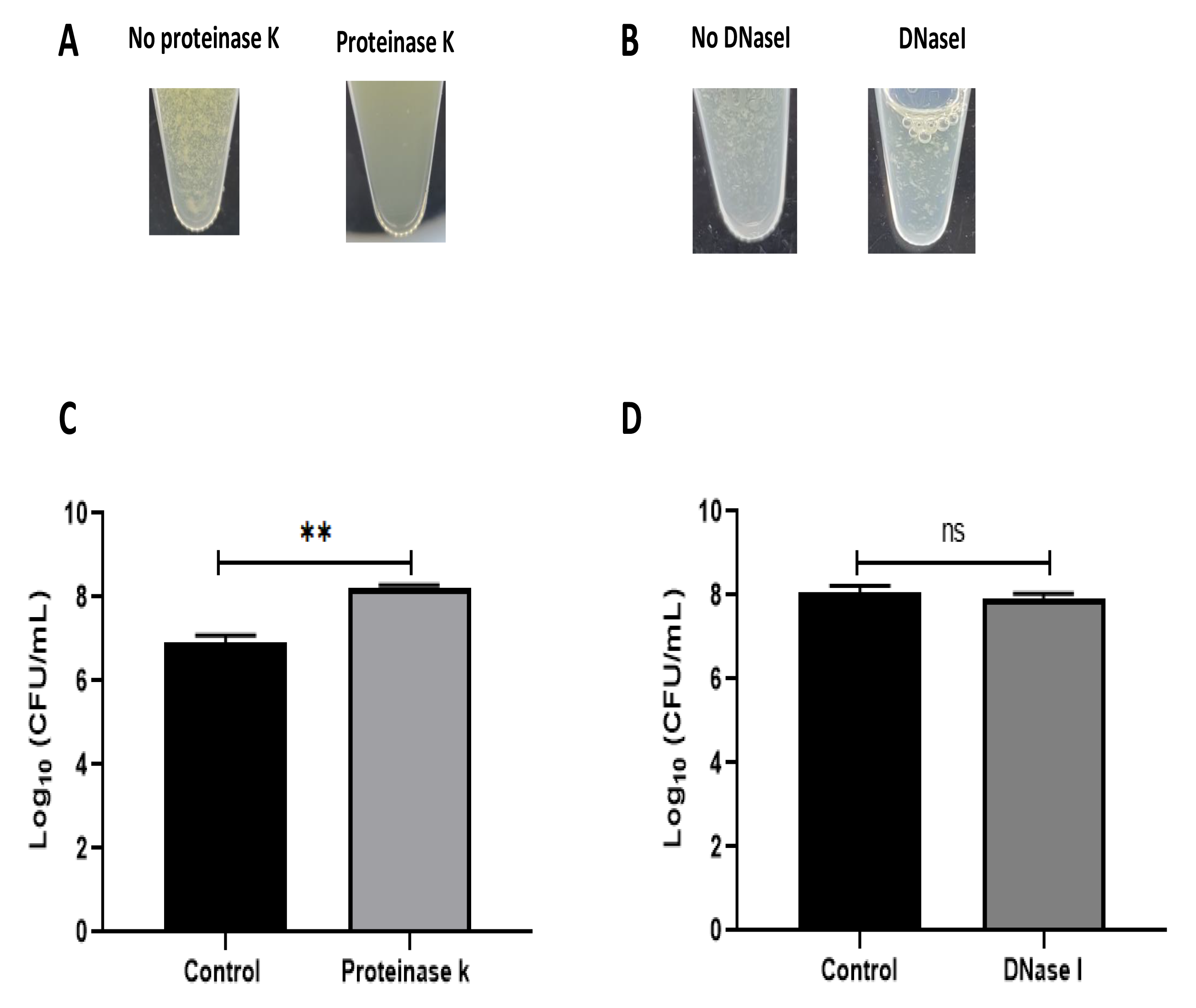
Biochemical disruption assay of *S. aureus* BP043 synovial fluid aggregates. **A)** Destruction of synovial fluid aggregates after adding proteinase K (150 µg/mL) for 1 hour at 37°C. **B)** No clear destruction of synovial fluid aggregates after adding DNase I (0.5 mg/mL dissolved in 2 mM MgCl_2_). **C)** Proteinase K treatment resulted in the release of more bacterial cells and increased the CFU compared to no treatment (1x PBS), mean ± SE, N = 4 pools of synovial fluids. **D)** DNase I treatment did not affect the bacterial load indicating the extracellular DNA is probably not a major component of the synovial fluid aggregates, ± SE, N = 3 pools of synovial fluids. Unpaired, two-tailed t-test was utilized to determine statistical difference, **p < 0.01, ns: non-significant.

**Figure 4.**
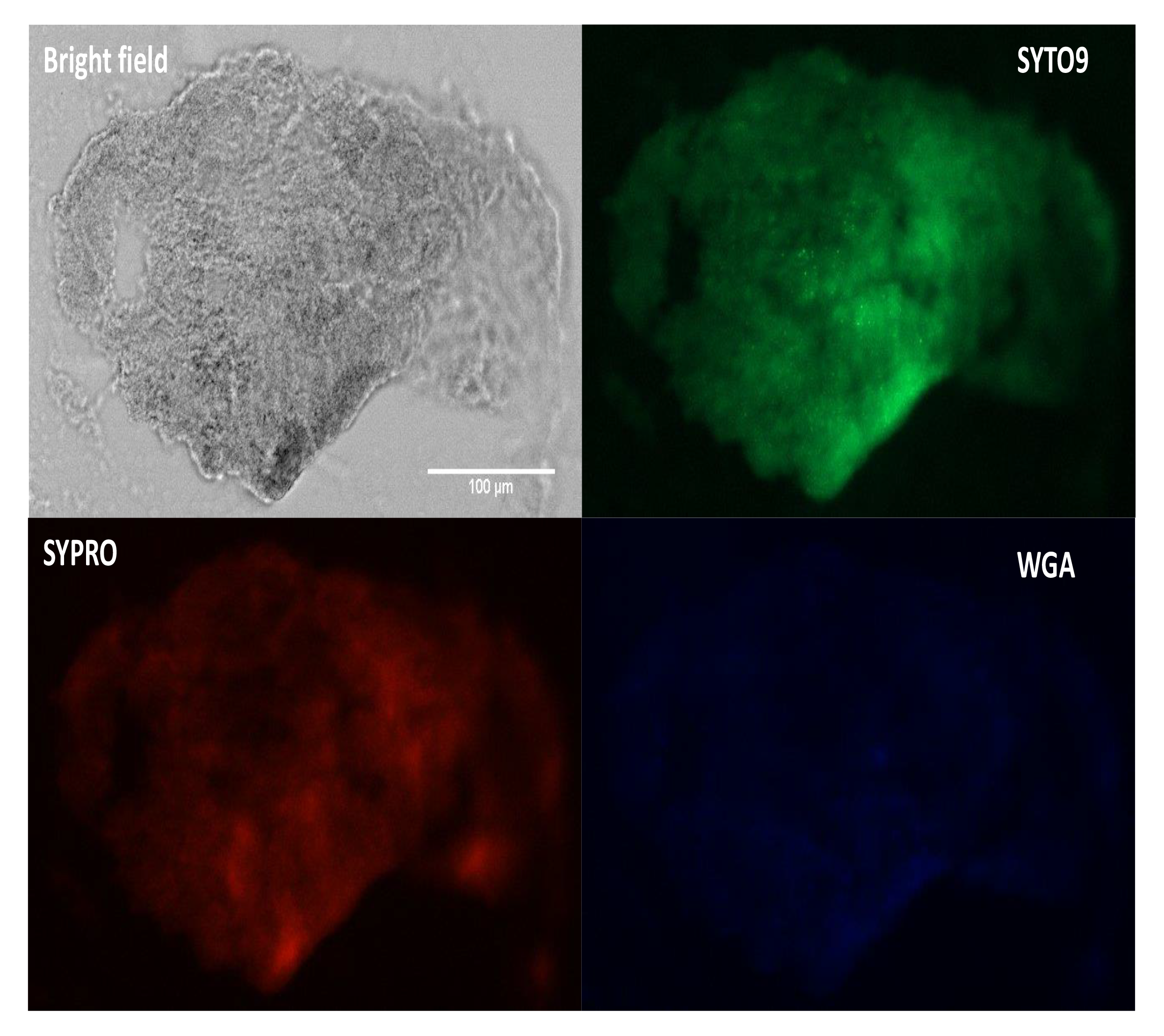
Fluorescence staining and microscopy examination of synovial fluid-derived *S. aureus* aggregates to detect the matrix composition. Aggregates were stained with SYTO9 (green) for nucleic acid (cellular biomass), SYPRO (red) for proteins and WGA (blue) for GlcNAc (carbohydrates). Scale bar = 100 µm.

### Phage Remus possess lytic activities against planktonic *S. aureus* BP043

To assess the lytic activity of phage Remus against planktonic *S. aureus* BP043, a virulence assay was performed. Figure 5 showed the dose-dependent effect of phage Remus against *S. aureus* BP043 planktonic culture for 48 hours at MOIs of 10 – 10^-6^. The lytic activities were most pronounced at higher MOIs of 10 – 10^-2^, where bacterial lysis occurred around 4 hours and was sustained for the next 44 hours. The optical densities were comparable to the blank (negative control) which were lacking bacteria (p > 0.05, no significant difference). At lower MOIs (10^-4^- 10^-6^), bacterial growth was detected at 48 hours and optical densities were significantly higher than the blank (p < 0.001). However, partial bacterial lysis was observed.

**Figure 5.**
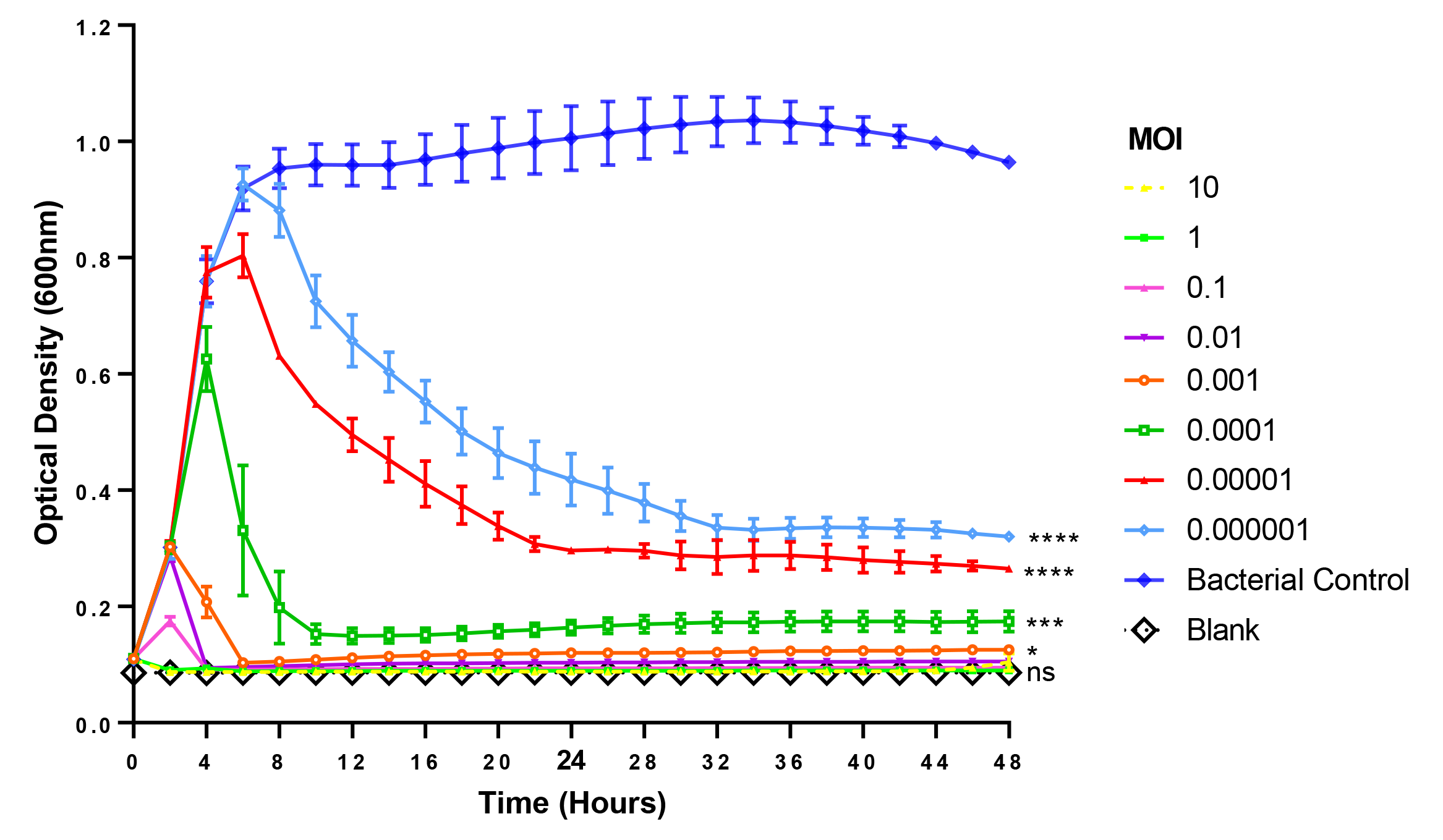
Lytic activities of phage Remus against *S. aureus* (BP043) planktonic cells (∼10^8^ CFU/mL) in TSB with Remus at 10-fold changes of MOIs ranging from 10-10^-6^. Phage Remus was applied right at the start of the experiment. Cell density was assessed by measuring optical densities at 600 nm every 120 minutes for 48 hours. N = 2, mean ± SE. One-way ANOVA (Tukey’s multiple comparisons test) was utilized to determine statistical difference at 48 hours. Asterisks show comparison between the different MOIs and the media blank (no bacteria). *P < 0.01, ***p < 0.001, ****p < 0.001, ns: non-significant.

### Phage Remus is effective against pre-formed *S. aureus* aggregates in synovial fluid

The lytic activities of Remus were assessed against pre-formed aggregates of *S. aureus* BP043 in synovial fluid. Several concentrations of Remus, ranging from 10^9^ to 10^6^ PFU/mL, were tested and *S. aureus* viability after 48 hours of incubation was compared across the different titers. Bacterial reduction was 8 log units at 10^9^ PFU/mL and decreased to 1.9 log at 10^6^ PFU/mL. The ability of Remus to significantly reduce bacterial load (p <0.0001) in aggregates at concentrations of 10^9^ to10^7^ PFU/mL versus the control samples without Remus treatment is demonstrated in Figure 6. Both concentrations, 10^9^ and 10^8^ PFU/mL, were the most effective in reducing bacterial viability in aggregates compared to 10^6^ PFU/mL (p < 0.0001) or 10^7^ PFU/mL (p = 0.0137, p = 0.0007, respectively).

**Figure 6.**
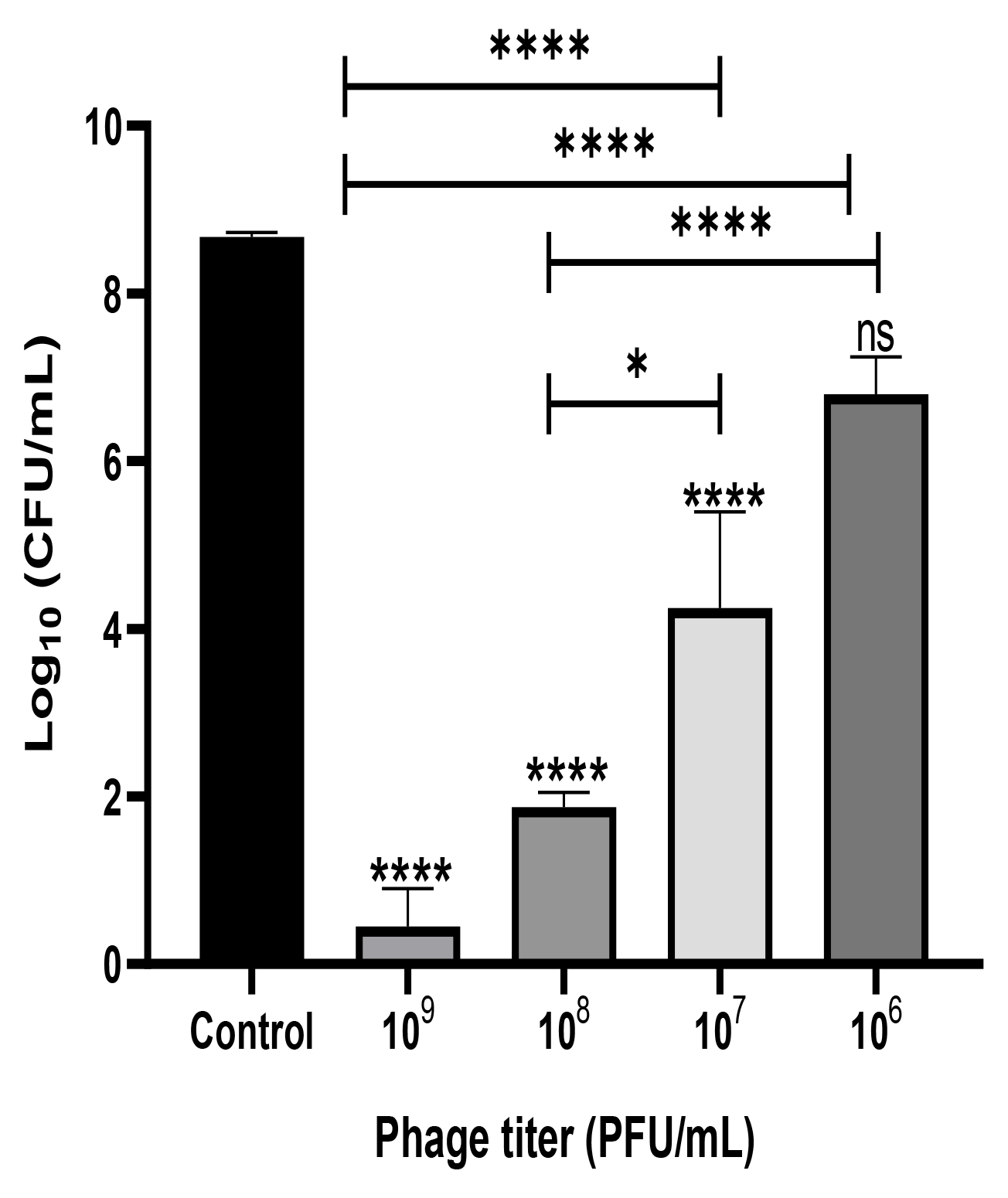
Phage Remus bacteriolytic activities against *S. aureus* BP043 preformed aggregates in human synovial fluid. Viable bacterial counts (log_10_ CFU/ml) after applying various titers of Remus (∼3×10^9^ - 10^6^ PFU/mL) for 48 hours against pre-formed aggregates of ∼4×10^8^ CFU/mL in synovial fluid (MOIs: 10 - 10^-2^). Control is non-phage treated aggregates. N = 2-6 synovial fluids, mean ± SE. Asterisks on top of the columns show comparison between the control (no treatment) and the different PFUs treatment groups. One-way ANOVA (Tukey’s multiple comparisons test) was utilized to determine statistical difference, *p < 0.05, ****p < 0.0001, ns: non-significant.

### Remus is capable of propagating in synovial fluid

We assessed phage survival and its ability to replicate in synovial fluid-derived aggregates after a 48 hour incubation period by comparing phage viability before and after incubation. A significant increase in Remus density (p= 0.005), from ∼ 1×10^8^ PFU/mL to 2×10^9^ PFU/mL was observed (Figure S1). This indicated that Remus could not only survive, but could propagate in an environment containing human synovial fluid.

### Combination therapy of phage and vancomycin is more effective than either agent alone *in vitro*

We tested whether phage Remus (10^8^ PFU/mL) had a better antimicrobial effect than vancomycin (500 µg/mL) against *S. aureus* biofilm-like aggregates in human synovial fluid. We also wanted to determine if vancomycin could interact synergistically with phage to treat these aggregates. Bacterial survival was monitored for the different treatment groups and was compared to the control. Treatment with phage Remus resulted in more than a 68% reduction in viable *S. aureus* residing in the synovial fluid aggregates, compared to the aggregates with no treatment (p < 0.0001, Figure 7). In figure 1, A we observed, macroscopically, the destruction of aggregates. Furthermore, Remus reduction of viable bacteria in BP043 aggregates was more pronounced compared to vancomycin-alone treatment which resulted in ∼ % 20 reduction in viability (p < 0.0001). Interestingly, treatment with phage Remus followed by vancomycin was more efficacious in reducing bacterial load (97% reduction) than using Remus or vancomycin alone (p = 0.0023, p< 0.0001, respectively). The coefficient of drug interaction (CDI) is equal to 0.0028 indicating significant synergistic interaction between the phage (Remus) and vancomycin.

**Figure 7.**
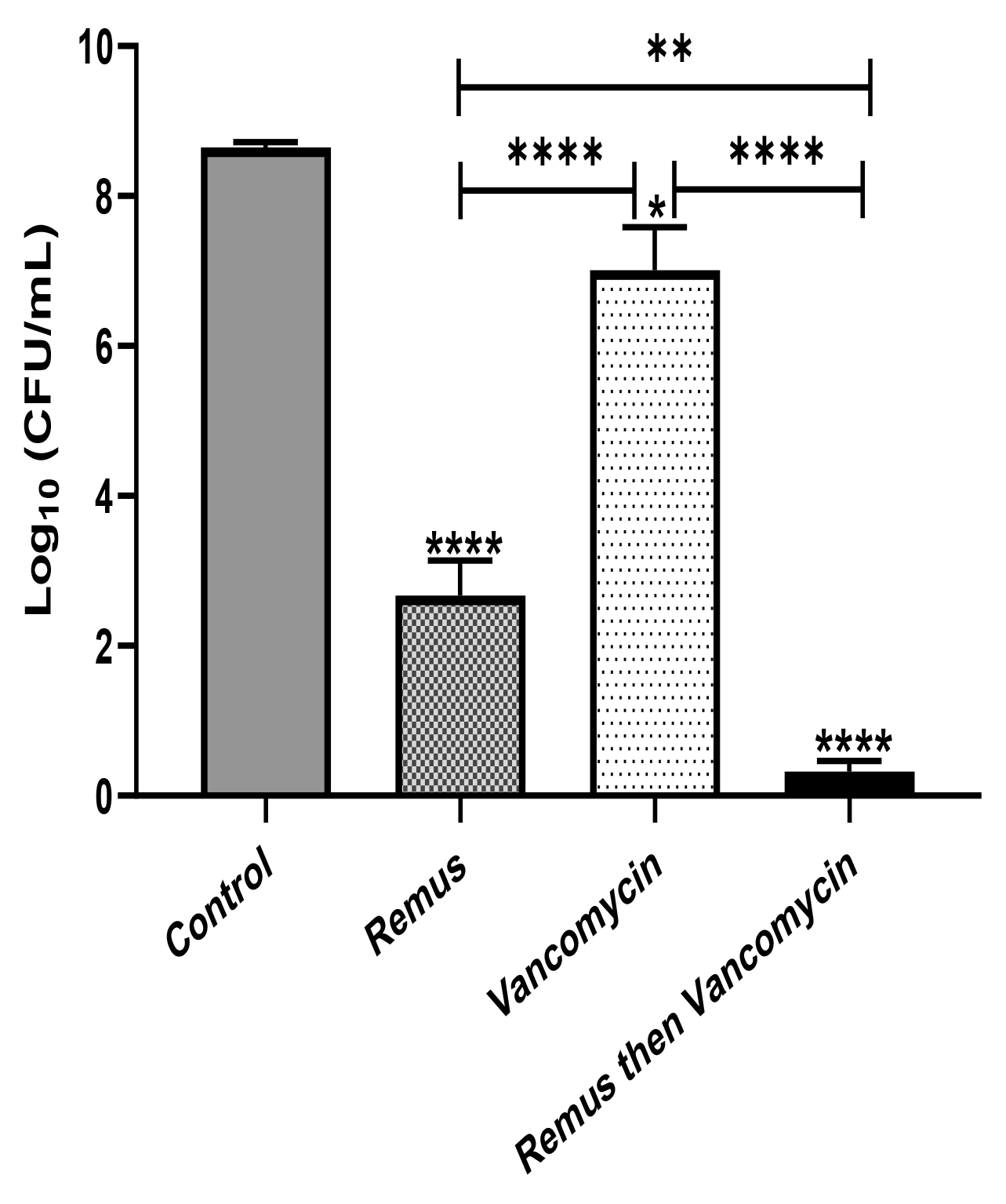
Phage Remus and vancomycin efficacy against pre-formed *S. aureus* BP043 aggregates in human synovial fluid. Bacterial load (log_10_ CFU/ml) in synovial fluid after applying phage Remus alone at ∼3×10^8^ PFU/mL (48 hours), vancomycin at 500 µg/mL (48 hours) and sequential treatment of phage (24 hours) followed by vancomycin (24 hours). N = 5 synovial fluids, mean ± SE. Asterisks on top of the columns show comparison between the control (no treatment) and the different treatment groups. Statistical significance was assessed by performing one-way ANOVA (Tukey’s multiple comparisons test), *p < 0.05, **p < 0.01, ****p < 0.0001.

### No development of Remus-resistance sub-population

The rise of a phage-resistant sub-population was monitored by determining the EOP of the *S. aureus* BP043 isolates that survived the 48 hour-Remus treatment in synovial fluid, compared to the ancestral strain *S. aureus* BP043. Figure S2 demonstrated no evolvement of resistance by *S. aureus* population that was exposed to Remus treatment (EOP= 1).

### The combination of phage Remus and vancomycin rescued *G. mellonella* larvae infected with *S. aureus* aggregates

Survival of *G. mellonella* larvae infected with *S. aureus* BP043 (as planktonic cells or synovial fluid-induced aggregates), was evaluated at 24-, 48-, 72- and 96- hours post-treatment with Remus and vancomycin alone or in combination (Figure 8, A and B).

**Figure 8.**
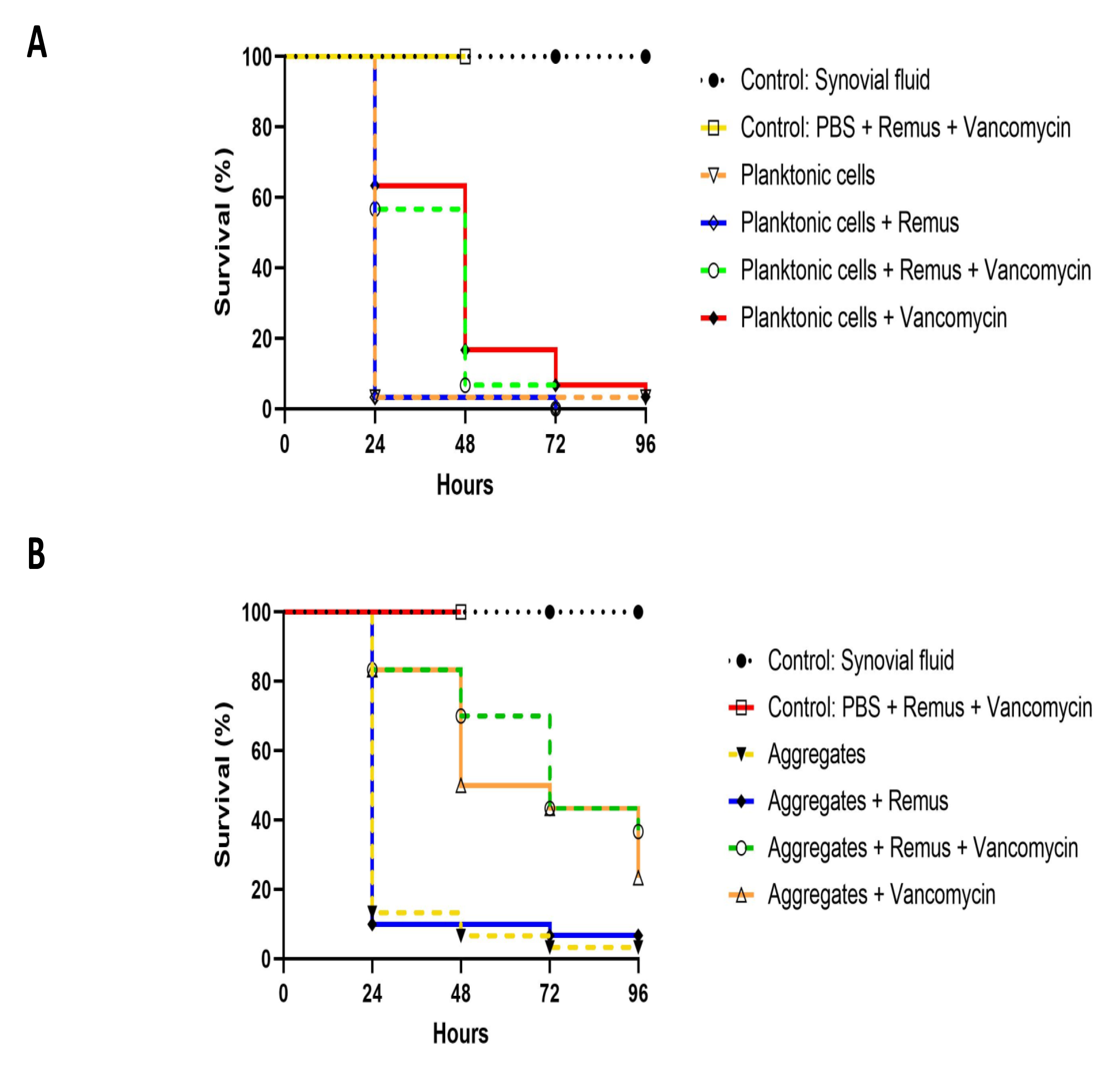
Kaplan-Meier survival analysis of *G. mellonella* infected with planktonic cells or synovial fluid aggregates of *S. aureus* BP043. Survival was evaluated after administration of 10^8^ PFU of phage Remus and/or 5 µg of vancomycin to larvae infected with *S. aureus* BP043 as, **A)** planktonic cells grown in Ringer’s solution TSB (2.2×10^7^ CFU) or **B)** aggregates (1.7×10^7^ CFU). Mortality was scored daily for 96 hours according to Table S1. Uninfected larvae receiving a Remus-vancomycin combination treatment were included as a control group up to 48 hours. The data represented are the mean of three biological replicates with ten larvae per group. Data of synovial fluid controls from the same experiments are presented in both figures for clarity and comparison. Statistically significant differences by the log-rank test were defined as p < 0.0001 between treatment groups.

#### Planktonic

The survival of untreated larvae infected with 2.2 x 10^7^ CFU of planktonic *S. aureus* decreased to 3% at 24 hours. Interestingly, these survival rates were less pronounced when compared to untreated larvae that were infected with aggregated *S. aureus* in which 13%, 7% and 3% survival was observed at 24-, 48- and 96- hours (Figure 8, A and B). Improved survival outcomes were observed at 24 hours post-treatment, where 63% and 57% survival were observed in groups receiving vancomycin and Remus-vancomycin treatments, respectively. This improvement was significant when compared to untreated larvae (3%; p < 0.0001). However, neither treatment was statistically significant when compared with each other (p = 0.60), but were significant when compared to treatment with Remus alone (p < 0.0001). Treatment with Remus alone did not rescue larvae (p > 0.9999). Additionally, comparison of survival curves over 48 hours revealed that treatment with either vancomycin alone (17%) or in combination with Remus (7%) continued to improve survival outcomes in comparison to untreated larvae (3%; p < 0.0001). Again, these treatments were not significantly different from each other (p = 0.31). Survival of infected larvae receiving Remus and vancomycin together, Remus alone or vancomycin alone resulted in 0%, 0% and 3%, respectively (Figure 8, A) which was comparable to the infected larvae with no treatment (3%) at 96 hours post-treatment. However, survival curves of infected larvae receiving either Remus-vancomycin or vancomycin alone were both significant when compared to untreated larvae (p < 0.001), whereas treatment with only Remus was not (p = 0.56). Moreover, the survival curve of the Remus-vancomycin treatment was not statistically significant compared to vancomycin alone (p = 0.22), but was more significant than treatment with Remus only (p < 0.0001). As anticipated, larvae in control groups that were either not injected or were injected with Remus and vancomycin did not display any signs of melanization or death.

#### Aggregates

A trend of improved survival rates over time in larva infected with *S. aureus* aggregates was observed with a phage-vancomycin treatment regime compared to phage or vancomycin, alone (Figure 8,B). At 24 hours post-treatment, 83% survival was observed in larvae receiving either vancomycin alone or in combination with Remus. This was a significant improvement when compared to larvae receiving no treatment (13%; p < 0.0001) or treatment with Remus alone (10%; p < 0.0001). Administration of Remus alone was not able to improve survival outcomes when compared to untreated larvae (p = 0.69). Moreover, analysis at 48 hours revealed that administration with vancomycin and the Remus-vancomycin combination treatment resulted in survival rates of 50% and 70%, respectively. These were significant when compared to untreated larvae (7%; p < 0.0001) and treatment with Remus alone (10%; p < 0.0001), but were not significant when compared to each other (p = 0.17). Survival in larvae receiving treatment with Remus alone was not significantly different from untreated larvae (p = 0.80). The administration of the Remus-vancomycin treatment against *S. aureus* aggregates resulted in the highest survival rate of larvae (37 %) at 96 hours post-treatment, compared to untreated larvae (3 %; p < 0.0001). Significant improvement was also observed in infected larvae receiving vancomycin only (23 %; p < 0.0001), however this was not the case larvae treated with phage only (7 %; p = 0.72), compared to infected untreated group. Survival with Remus-vancomycin treatment was not statistically more significant when compared to administration of vancomycin alone (23 %; p = 0.31), but was more significant in comparison to phage-only treatment group (7 %; p < 0.0001), 96 hours post-treatment (Figure 8,B).

### Bioinformatic analysis of Remus putative tail fibers

The morphogenesis module of the Remus genome was examined for genes encoding tail-associated proteins. Of the 23 candidate proteins, only ten were returned with database hits (Table S2). Of the ten proteins with functional domains, gene products (gp) 134, and 137 were excluded from further analysis due to the hits being domains of unknown function. The gp138 and 140 proteins contained hits to baseplate structural proteins and no enzymatic hits, suggesting they may not play a role in biofilm depolymerization and were thus not studied further (Table S2).

Six proteins of interest, gp141, and 144 – 148, were chosen for further study with PHYRE2 because they all feature InterProScan functional domain hits associated with teichoic acid, cystine, or peptidoglycan hydrolases: phospholipase C (PLC)-like phosphodiesterase (gp144), CHAP proteinase (cysteine, histidine-dependent amidohydrolases/peptidases) (gp145 and 146), and lysozyme domains (gp141, 147 and 148) (Table S2). PHYRE2 analysis of each protein supports the InterProScan findings (gp141 and gp144-148). Gp141 was modelled to a T4 gp25 tail lysozyme (99.8% confidence and 19% identity). It should be noted, gp141 is an interesting protein as it does contain a lysozyme domain, but investigation into other proteins with this domain architecture revealed they showed no lysozyme activity (41). This suggests while gp141 encodes a lysozyme domain, it may not be used as a main driver of peptidoglycan depolymerization. The top PHYRE2 model of gp144 is to *Bacillus subtilis* 168 GlpQ, a phosphate starvation-induced wall teichoic acid (WTA) hydrolase with 100% confidence and 25% identity (gp144). Both gp145 and gp146 proteins modelled to CHAP-domain containing proteins, with the gp145 top PHYRE2 hit to γ-D-glutamyl-L-diamino endopeptidase from *Nostoc punctiforme* PCC 73102 (100% confidence; 18% identity), and gp146 modeling to n-acetylmuramoyl-l-alanine amidase domain-containing protein of *S. aureus* (100% confidence; 26 % identity) (gp145-146). Finally, both gp147 and 148 PHYRE2 models are to proteins with known lysozyme activity: the gp147 model corresponds to a cell wall degrading enzyme from *B. subtilis* phage phi29 (100% confidence; 20% identity), while the gp148 protein models to the glucosaminidase domain of the bifunctional autolysin AltA from *S. aureus* (100% confidence; 29% identity) (gp147-148).

## Discussion

The aim of this study was to evaluate if phages have a better antimicrobial effect than vancomycin against *S. aureus* biofilm-like-aggregates formed during incubation in human synovial fluid.

Several studies reported the ability of staphylococci to form free-floating, biofilm-like aggregates in human and animal synovial fluid *in vitro* (9-12, 37). Similarly, infected synovial fluid samples collected from patients can also accommodate aggregates (10). In the current work, we have shown the same phenotype of biofilm-like aggregates in 78% human synovial fluid by a PJI clinical isolate of *S. aureus* BP043 (MRSA). The SEM images demonstrated the presence of fibrinogen-like filaments as part of these aggregates. Previous studies had identified the presence of fibrinogen in synovial fluid and fibronectin-and fibrinogen-binding proteins existing on the cell wall of *S. aureus* as important factors in the process of aggregates formation, thereby supporting our observations (10, 11, 42). It has been shown previously that *S. aureus* can form more robust aggregates compared to other staphylococci, such as *S. epidermidis* and *S. lugdunesis* (10-12, 42). It is thought that these biofilm-like aggregates negatively impact PJI treatments, due to an increased ability to resist antibiotic treatment (10, 13). Bidossi *et al*. reported that the MIC for the tested *S. aureus* strains against vancomycin and rifampin was 4 to 32 times higher in synovial fluid compared to the MIC obtained in Muller Hinton broth (MH)(10). The same pattern was observed for pre-formed aggregates (10). Moreover, this group showed that the enzymatic treatment of the pre-formed aggregates increased their susceptibility to antibiotic treatment. *S. aureus* BP043 strain has a MIC of 2 µg/mL for vancomycin in MH(19). However, when BP043 biofilm-like aggregates in synovial fluid were exposed to vancomycin at 250 times higher MIC (500 µg/mL), the bacterial cells survived the treatment. Putting all of this together, we proposed that the biofilm-like, aggregative phenotype in synovial fluid offers protection and increases vancomycin tolerance (42). This has motivated the use of phage as a potential or adjunct treatment strategy to antibiotics.

We examined the efficacy of phage therapy against these human-derived synovial fluid aggregates of *S. aureus*. This study is the first to test phage treatment against synovial fluid-induced aggregates *in vitro* and *in vivo.* Some phages, and particularly Remus, had been reported to have antibiofilm activities (20). We have shown that Remus is capable of decomposing aggregate biomass and reducing bacterial load *in vitro* in a dose-dependent manner. This makes Remus a potentially promising candidate for therapeutic applications, especially since it possesses remarkable and rapid lytic activities against planktonic cells. Phages contain genes that encode enzymes such as endolysins, virion-associated peptidoglycan hydrolases (VAPGHs) and polysaccharide depolymerases which are capable of disrupting various biofilm components, such as peptidoglycan, polysaccharides, and proteins (17). Bioinformatic analysis of the Remus genome suggested that there are 6 putative lysin genes in the tail morphogenesis region: a WTA hydrolase, two CHAP peptidases, and two lysozymes. Each of these proteins are functionally-relevant enzymes that could contribute to biofilm disruption capabilities, as seen with other phage-derived lysins. Purified lysins, or peptidoglycan hydrolases, have bactericidal effects on susceptible bacteria by degrading the bacterial cell wall resulting in cell lysis. There are four distinct activities of peptidoglycan hydrolases: cleaving the peptidoglycan between the sugar moieties, such as endo-β-N-acetylglucosaminidase or N-acetylmuramidase, cleavage between the stem peptide and sugar moieties (N-acetylmuramoyl-L-alanine amidase), or between amino acids in the stem peptide or cross bridge (endopeptidases) (43). Significant research has been conducted on the use of lysins for the eradication of antimicrobial resistant Gram-positive infections and biofilms (44-47). Fenton *et al.* reported the capacity of the purified lysin CHAP to completely remove established *S. aureus* biofilm and lyse bacterial cells. It was suggested that the lysing activity of CHAP could be responsible for biofilm destabilization and detachment rather than acting directly on the biofilm matrix components (48). A similar pattern of quick biofilm removal as a result of lysing bacterial cells within a biofilm was seen with SAL-2, acell-wall-degrading enzyme (14).

Sass *et al.* explored the activities of ϕ11 lysin against *S. aureus*. This lysin possesses three active domains; CHAP domain, amidase domain, and a cell wall binding domain, which had been reported to play a role in ϕ11’s ability to lyse the bacterial cells and disrupt of staphylococcus biofilm (49). The team reported both antibacterial and antibiofilm activities. Interestingly, ϕ11 lysin was not capable of removing the polysaccharides biofilm matrix of *S. epidermidis* O-47, but could degrade the *S. aureus* NCTC8325 proteinaceous and polysaccharides biofilm matrix. Therefore, it was suggested that ϕ11 lysin is capable of acting against the protein part of the biofilm matrix of *S. aureus*. Another lysin that is potentially produced by Remus is the WTA hydrolase. It works by targeting the cell wall teichoic acid. Teichoic acid is a carbohydrate-containing polymer found in the cell wall of all Gram-positive bacteria (27). WTAs contribute to bacterial cell adhesion, the first step of biofilm formation, to biotic and abiotic surfaces (50-52). Evidence suggests that teichoic acid increases bacterial attachment to fibronectin-coated surfaces (51, 53). Absence or degradation of WTAs by WTA hydrolases affects the initial attachment and biofilm forming ability of the bacteria (50, 54). Interestingly, it had been shown that some species such as *S. epidermidis* produces extracellular teichoic acid that will be part of the biofilm matrix (55).

In the current study, Remus had shown its capacity to eradicate the synovial fluid-derived aggregates and lyse the embedded *S. aureus* cells. Our data demonstrated the macroscopic biomass degradation and the reduction in the viable bacterial count. This could be attributed to Remus’ ability to lyse bacterial cells through its lysins (CHAP, wall teichoic acid hydrolase and lysozymes) resulting in the destabilization and destruction of the aggregates. The biochemical disruption assay and fluorescence microscopy data, in this current work, and in other groups’ reports indicated the same observation of a proteinaceous nature of the aggregates (10, 42). This suggests that Remus might have some direct enzymatic activities against the protein components of the aggregate matrix. However, the only evidence that we currently have is that Remus’ lysins function as peptidoglycan hydrolases. More evidence and experiments should be conducted to show the specific enzymatic activities of Remus and any potential direct degradation power of the lysins against the aggregate matrix chemical components.

We investigated the efficacy of the combination therapy of phage Remus and vancomycin compared to using either agent alone *in vitro*. We observed better clearance of synovial fluid aggregates of *S. aureus* MRSA isolate when both Remus and vancomycin were used together and applied sequentially compared to using Remus or vancomycin separately. They interacted synergistically to reduce the viability of bacterial cells hiding in robust human synovial fluid aggregates. Previously (56), our group and others showed a similar pattern of synergistic interaction between the phages and antibiotics, leading to the highest reduction against broth-derived biofilm compared to using single agents (35, 56-58). The phage-antibiotic interaction was mainly noted at sub-lethal doses of antibiotics (35, 56, 58). Some reported that the efficiency of the combination treatment was significantly pronounced when the two agents were applied separately, specifically when phage was used prior to the antibiotic compared to simultaneous application (35, 56,58, 59).

It was proposed that the treatment of biofilms with phage preceding the antibiotic allows phages to rapidly replicate in the bacterially-dense environment of the biofilm in regular lab media. This would result in high phage densities and the disruption of the biofilm matrix. A number of factors could affect the outcome of phage and antibiotic interaction, such as the type of tested bacteria, phage and antibiotic. These data drove the usage of the sequential treatment strategy for phage Remus and vancomycin against *S. aureus* synovial fluid-induced aggregates in the current study.

*G. mellonella* larvae were used to assess the *in vivo* application of Remus and vancomycin alone or together to treat *S. aureus* infections with planktonic cells or synovial fluid induced aggregates. The combination treatment of phage-vancomycin was effective in rescuing 37% of *G. mellonella* larvae infected with *S. aureus* aggregates after 96 hours post-treatment, respectively. Additionally, administration of a combination therapy significantly improved daily survival outcomes after 24 and 48 hours post-treatment, resulting in 83% and 70% survival, respectively. These survival rates were not achievable when phage-alone or vancomycin-alone were administered. At the 96-hour time point, there is a clear distinction between the uninfected larvae (100% survival) and the untreated larvae (nearly 0% survival), so any treatment (vancomycin, phage or the combination) that improves survival from 0% implies treatment success. Given the lethality of the inoculum, a complete reversal of mortality would be a challenging prospect to expect and offers too high of a bar to overcome. Thus, since phage and vancomycin together resulted in the highest survival of any of the treatments (37% survival versus 23% for vancomycin alone and 7% for phage alone) after 96 hours, this is remarkable from the untreated group (3%). Moreover, within the treatment groups, the 14% increase in survivability demonstrated with phage and vancomycin combined, suggest synergy and matches our trend from the *in vitro* data.

In contrast, the combination and phage-alone treatments were not effective in rescuing larvae infected with planktonic bacteria after 96 hours post-treatment, even though our *in vitro* data showed efficiency at lower phage titers (10^8^ −10^6^ PFU/mL, MOI= 0.01, 0.1 and 1). This could be attributed to the fact that before treatment administration and during the 1- hour incubation post-injection with planktonic cells, irreversible toxicity to the larvae occurred. This could be due to high bacterial dose (2×10^7^ CFU) and induced virulence. Our data showed that when larvae were infected with a lower bacterial dose of planktonic cells of BP043 (4.1×10^6^ CFU), bacterial survival was 40% at 24 hours and 30% at 48 hours (Data not shown). However, rescue of larvae infected with planktonic bacteria was shown to be most significant at 24 hours with administration of vancomycin (63%) or the combination therapy (57%). Interestingly, BP043 aggregates at a similar concentration (1×10^7^ CFU) of planktonic cells (2×10^7^ CFU) showed better survival rates of the larvae. The observed difference in virulence between planktonic and aggregate bacterial population could be due to the possibility that bacterial cells present within aggregates are in a more dormant form.

Limitations of this study include: investigating a single strain of *S. aureus* as well as testing the efficacy of a single phage agent. However, there are numerous strengths of this study including the use of a clinically-relevant, PJI-isolated, *S. aureus* strain and testing it in clinical samples of synovial fluid. In addition, this is the first study to test the efficacy of Remus against *S. aureus* biofilm-like aggregates formed in synovial fluid. The *G. mellonella* assay has also shown to be a valuable tool in translating our findings into larger animal models that are more clinically representative of PJI.

Our data indicates that targeting synovial fluid-derived, biofilm-like aggregates with a combination of phage Remus and vancomycin resulted in synergistic interaction and a notable clearance of the infection. This work is aimed at gathering preclinical evidence for using phage as a new therapeutic avenue to treat PJI infections.

## Contribution to the field statement

Prosthetic joint infection (PJI) is a devastating complication that can occur following joint replacement surgeries. Current therapies for PJI have a failure rate reaching 30%. Treatment failure can lead to limb amputations and death. Biofilm formation is a major clinical challenge contributing to antibiotic tolerance and treatment failure of PJI. Biofilms are characterized as interconnected communities of microorganisms that adhere on to surfaces and interconnect within a self-produced matrix that protects bacterial cells from antibiotics and the host immune system. Recent studies demonstrate that staphylococci are capable of forming biofilm-like clumps in synovial fluid. These biofilm-like aggregates elevate the minimum inhibitory concentration of antibiotics necessary to control an infection. Lytic bacteriophages (phages) can target biofilm-associated bacteria at localized sites of infection. The aim of this study was to test if phage and vancomycin are capable of clearing S*taphylococcus aureus* (the most isolated pathogen from PJI patients) formed in synovial fluid.

This work demonstrated, for the first time, the efficiency of phage treatment against MRSA biofilm-like aggregates. Moreover, this efficiency is augmented when phage Remus and vancomycin are combined leading to better clearance of MRSA synovial fluid aggregates. This work is aimed at gathering preclinical evidence for using phage as therapeutic avenue to treat PJI.

Figure S1. Phage Remus titer in synovial fluid-derived aggregates of *S. aureus* BP043 aggregates. Remus density was checked after 48 hours at 37°C and compared to initial titer. N = 4, ± SE. Statistical significance was performed using t-test (two-tailed, unpaired). **p<0.01.

Figure S2. No development of resistance by *S. aureus* BP043 against Remus. The rise of phage-resistance sub-population was monitored by checking the efficiency of plating (EOP) of the *S. aureus* BP043 isolates that survived the 48 hours Remus treatment or for the control (no Remus treatment) in synovial fluid (SF). EOP was calculated by dividing Remus titer on the tested *S. aureus* BP043 by Remus titer on the ancestor *S. aureus* BP043. Two synovial fluids were used (SF1, SF2), and four *S. aureus* isolates were checked for resistance per synovia fluid.

## References

1. Kunutsor SK, Whitehouse MR, Blom AW, Board T, Kay P, Wroblewski BM, et al. One-and two-stage surgical revision of peri-prosthetic joint infection of the hip: a pooled individual participant data analysis of 44 cohort studies. Eur J Epidemiol. 2018;33(10):933–46.

2. Hartzler MA, Li K, Geary MB, Odum SM, Springer BD. Complications in the treatment of prosthetic joint infection. Bone Joint J. 2020;102-B(6_Supple_A):145-50.

3. Lenguerrand E, Whitehouse MR, Beswick AD, Toms AD, Porter ML, Blom AW, et al. Description of the rates, trends and surgical burden associated with revision for prosthetic joint infection following primary and revision knee replacements in England and Wales: an analysis of the National Joint Registry for England, Wales, Northern Ireland and the Isle of Man. BMJ Open. 2017;7(7):e014056.

4. Gunther F, Wabnitz GH, Stroh P, Prior B, Obst U, Samstag Y, et al. Host defence against Staphylococcus aureus biofilms infection: phagocytosis of biofilms by polymorphonuclear neutrophils (PMN). Mol Immunol. 2009;46(8-9):1805–13.

5. Singh R, Ray P, Das A, Sharma M. Penetration of antibiotics through Staphylococcus aureus and Staphylococcus epidermidis biofilms. J Antimicrob Chemother. 2010;65(9):1955–8.

6. Thurlow LR, Hanke ML, Fritz T, Angle A, Aldrich A, Williams SH, et al. Staphylococcus aureus biofilms prevent macrophage phagocytosis and attenuate inflammation in vivo. J Immunol. 2011;186(11):6585–96.

7. Tande AJ, Patel R. Prosthetic joint infection. Clin Microbiol Rev. 2014;27(2):302–45.

8. Stoodley P, Nistico L, Johnson S, Lasko LA, Baratz M, Gahlot V, et al. Direct demonstration of viable Staphylococcus aureus biofilms in an infected total joint arthroplasty. A case report. J Bone Joint Surg Am. 2008;90(8):1751–8.

9. Staats A, Burback PW, Eltobgy M, Parker DM, Amer AO, Wozniak DJ, et al. Synovial Fluid-Induced Aggregation Occurs across Staphylococcus aureus Clinical Isolates and is Mechanistically Independent of Attached Biofilm Formation. Microbiol Spectr. 2021;9(2):e0026721.

10. Bidossi A, Bottagisio M, Savadori P, De Vecchi E. Identification and Characterization of Planktonic Biofilm-Like Aggregates in Infected Synovial Fluids From Joint Infections. Front Microbiol. 2020;11:1368.

11. Knott S, Curry D, Zhao N, Metgud P, Dastgheyb SS, Purtill C, et al. Staphylococcus aureus Floating Biofilm Formation and Phenotype in Synovial Fluid Depends on Albumin, Fibrinogen, and Hyaluronic Acid. Front Microbiol. 2021;12:655873.

12. Perez K, Patel R. Biofilm-like aggregation of Staphylococcus epidermidis in synovial fluid. J Infect Dis. 2015;212(2):335–6.

13. Gilbertie JM, Schnabel LV, Hickok NJ, Jacob ME, Conlon BP, Shapiro IM, et al. Equine or porcine synovial fluid as a novel ex vivo model for the study of bacterial free-floating biofilms that form in human joint infections. PLoS One. 2019;14(8):e0221012.

14. Son JS, Lee SJ, Jun SY, Yoon SJ, Kang SH, Paik HR, et al. Antibacterial and biofilm removal activity of a podoviridae Staphylococcus aureus bacteriophage SAP-2 and a derived recombinant cell-wall-degrading enzyme. Appl Microbiol Biotechnol. 2010;86(5):1439–49.

15. Jassim SA, Limoges RG. Natural solution to antibiotic resistance: bacteriophages ’The Living Drugs’. World J Microbiol Biotechnol. 2014;30(8):2153–70.

16. Briandet R, Lacroix-Gueu P, Renault M, Lecart S, Meylheuc T, Bidnenko E, et al. Fluorescence correlation spectroscopy to study diffusion and reaction of bacteriophages inside biofilms. Appl Environ Microbiol. 2008;74(7):2135–43.

17. Chan BK, Abedon ST. Bacteriophages and their enzymes in biofilm control. Curr Pharm Des. 2015;21(1):85–99.

18. Broudy TB, Pancholi V, Fischetti VA. The in vitro interaction of Streptococcus pyogenes with human pharyngeal cells induces a phage-encoded extracellular DNase. Infect Immun. 2002;70(6):2805–11.

19. Taha M, Arulanandam R, Chen A, Diallo JS, Abdelbary H. Combining povidone-iodine with vancomycin can be beneficial in reducing early biofilm formation of methicillin-resistant Staphylococcus aureus and methicillin-sensitive S. aureus on titanium surface. J Biomed Mater Res B Appl Biomater. 2023;111(5):1133–41.

20. Vandersteegen K, Kropinski AM, Nash JH, Noben JP, Hermans K, Lavigne R. Romulus and Remus, two phage isolates representing a distinct clade within the Twortlikevirus genus, display suitable properties for phage therapy applications. J Virol. 2013;87(6):3237–47.

21. Aida Y, Pabst MJ. Removal of endotoxin from protein solutions by phase separation using Triton X-114. J Immunol Methods. 1990;132(2):191–5.

22. Mazzocco A, Waddell TE, Lingohr E, Johnson RP. Enumeration of bacteriophages using the small drop plaque assay system. Methods Mol Biol. 2009;501:81–5.

23. Gutierrez D, Ruas-Madiedo P, Martinez B, Rodriguez A, Garcia P. Effective removal of staphylococcal biofilms by the endolysin LysH5. PLoS One. 2014;9(9):e107307.

24. Loza-Correa M, Ayala JA, Perelman I, Hubbard K, Kalab M, Yi QL, et al. The peptidoglycan and biofilm matrix of Staphylococcus epidermidis undergo structural changes when exposed to human platelets. PLoS One. 2019;14(1):e0211132.

25. Wright CS. Structural comparison of the two distinct sugar binding sites in wheat germ agglutinin isolectin II. J Mol Biol. 1984;178(1):91–104.

26. Chen JK, Shen CR, Liu CL. N-acetylglucosamine: production and applications. Mar Drugs. 2010;8(9):2493–516.

27. Otto M. Staphylococcal biofilms. Curr Top Microbiol Immunol. 2008;322:207–28.

28. Konopka JB. N-acetylglucosamine (GlcNAc) functions in cell signaling. Scientifica (Cairo). 2012;20121.

29. van Dalen R, Peschel A, van Sorge NM. Wall Teichoic Acid in Staphylococcus aureus Host Interaction. Trends Microbiol. 2020;28(12):985–98.

30. Berggren K, Chernokalskaya E, Steinberg TH, Kemper C, Lopez MF, Diwu Z, et al. Background-free, high sensitivity staining of proteins in one-and two-dimensional sodium dodecyl sulfate-polyacrylamide gels using a luminescent ruthenium complex. Electrophoresis. 2000;21(12):2509–21.

31. Liu C, Bayer A, Cosgrove SE, Daum RS, Fridkin SK, Gorwitz RJ, et al. Clinical practice guidelines by the infectious diseases society of america for the treatment of methicillin-resistant Staphylococcus aureus infections in adults and children: executive summary. Clin Infect Dis. 2011;52(3):285–92.

32. Davis JS, Van Hal S, Tong SY. Combination antibiotic treatment of serious methicillin-resistant Staphylococcus aureus infections. Semin Respir Crit Care Med. 2015;36(1):3–16.

33. Feng Y, Tonon CC, Hasan T. Dramatic destruction of methicillin-resistant Staphylococcus aureus infections with a simple combination of amoxicillin and light-activated methylene blue. J Photochem Photobiol B. 2022;235:112563.

34. Xu SP, Sun GP, Shen YX, Peng WR, Wang H, Wei W. Synergistic effect of combining paeonol and cisplatin on apoptotic induction of human hepatoma cell lines. Acta Pharmacol Sin. 2007;28(6):869–78.

35. Chaudhry WN, Concepcion-Acevedo J, Park T, Andleeb S, Bull JJ, Levin BR. Synergy and Order Effects of Antibiotics and Phages in Killing Pseudomonas aeruginosa Biofilms. PLoS One. 2017;12(1):e0168615.

36. Wright RCT, Friman VP, Smith MCM, Brockhurst MA. Resistance Evolution against Phage Combinations Depends on the Timing and Order of Exposure. mBio. 2019;10(5).

37. Pestrak MJ, Gupta TT, Dusane DH, Guzior DV, Staats A, Harro J, et al. Investigation of synovial fluid induced Staphylococcus aureus aggregate development and its impact on surface attachment and biofilm formation. PLoS One. 2020;15(4):e0231791.

38. Loh JM, Adenwalla N, Wiles S, Proft T. Galleria mellonella larvae as an infection model for group A streptococcus. Virulence. 2013;4(5):419–28.

39. Blum M, Chang HY, Chuguransky S, Grego T, Kandasaamy S, Mitchell A, et al. The InterPro protein families and domains database: 20 years on. Nucleic Acids Res. 2021;49(D1):D344–D54.

40. Jones P, Binns D, Chang HY, Fraser M, Li W, McAnulla C, et al. InterProScan 5: genome-scale protein function classification. Bioinformatics. 2014;30(9):1236–40.

41. Lossi NS, Dajani R, Freemont P, Filloux A. Structure-function analysis of HsiF, a gp25- like component of the type VI secretion system, in Pseudomonas aeruginosa. Microbiology (Reading). 2011;157(Pt 12):3292–305.

42. Dastgheyb S, Parvizi J, Shapiro IM, Hickok NJ, Otto M. Effect of biofilms on recalcitrance of staphylococcal joint infection to antibiotic treatment. J Infect Dis. 2015;211(4):641–50.

43. Lood R, Molina H, Fischetti VA. Determining bacteriophage endopeptidase activity using either fluorophore-quencher labeled peptides combined with liquid chromatography-mass spectrometry (LC-MS) or Forster resonance energy transfer (FRET) assays. PLoS One. 2017;12(3):e0173919.

44. Vazquez R, Garcia E, Garcia P. Phage Lysins for Fighting Bacterial Respiratory Infections: A New Generation of Antimicrobials. Front Immunol. 2018;9:2252.

45. Gilmer DB, Schmitz JE, Euler CW, Fischetti VA. Novel bacteriophage lysin with broad lytic activity protects against mixed infection by Streptococcus pyogenes and methicillin-resistant Staphylococcus aureus. Antimicrob Agents Chemother. 2013;57(6):2743–50.

46. Sharma U, Vipra A, Channabasappa S. Phage-derived lysins as potential agents for eradicating biofilms and persisters. Drug Discov Today. 2018;23(4):848–56.

47. Ferriol-Gonzalez C, Domingo-Calap P. Phages for Biofilm Removal. Antibiotics (Basel). 2020;9(5).

48. Fenton M, Keary R, McAuliffe O, Ross RP, O’Mahony J, Coffey A. Bacteriophage-Derived Peptidase CHAP(K) Eliminates and Prevents Staphylococcal Biofilms. Int J Microbiol. 2013;2013:625341.

49. Sass P, Bierbaum G. Lytic activity of recombinant bacteriophage phi11 and phi12 endolysins on whole cells and biofilms of Staphylococcus aureus. Appl Environ Microbiol. 2007;73(1):347–52.

50. Vergara-Irigaray M, Maira-Litran T, Merino N, Pier GB, Penades JR, Lasa I. Wall teichoic acids are dispensable for anchoring the PNAG exopolysaccharide to the Staphylococcus aureus cell surface. Microbiology (Reading). 2008;154(Pt 3):865–77.

51. Hussain M, Heilmann C, Peters G, Herrmann M. Teichoic acid enhances adhesion of Staphylococcus epidermidis to immobilized fibronectin. Microb Pathog. 2001;31(6):261–70.

52. Aly R, Shinefield HR, Litz C, Maibach HI. Role of teichoic acid in the binding of Staphylococcus aureus to nasal epithelial cells. J Infect Dis. 1980;141(4):463–5.

53. Aly R, Levit S. Adherence of Staphylococcus aureus to squamous epithelium: role of fibronectin and teichoic acid. Rev Infect Dis. 1987;9 Suppl 4:S341–50.

54. Hou W, Kang S, Chang J, Tian X, Shi C. Correlation Analysis between GlpQ-Regulated Degradation of Wall Teichoic Acid and Biofilm Formation Triggered by Lactobionic Acid in Staphylococcus aureus. Foods. 2022;11(21).

55. Hussain M, Wilcox MH, White PJ. The slime of coagulase-negative staphylococci: biochemistry and relation to adherence. FEMS Microbiol Rev. 1993;10(3-4):191–207.

56. Kumaran D, Taha M, Yi Q, Ramirez-Arcos S, Diallo JS, Carli A, et al. Does Treatment Order Matter? Investigating the Ability of Bacteriophage to Augment Antibiotic Activity against Staphylococcus aureus Biofilms. Front Microbiol. 2018;9:127.

57. Cha Y, Chun J, Son B, Ryu S. Characterization and Genome Analysis of Staphylococcus aureus Podovirus CSA13 and Its Anti-Biofilm Capacity. Viruses. 2019;11(1).

58. Dickey J, Perrot V. Adjunct phage treatment enhances the effectiveness of low antibiotic concentration against Staphylococcus aureus biofilms in vitro. PLoS One. 2019;14(1):e0209390.

59. Tkhilaishvili T, Wang L, Perka C, Trampuz A, Gonzalez Moreno M. Using Bacteriophages as a Trojan Horse to the Killing of Dual-Species Biofilm Formed by Pseudomonas aeruginosa and Methicillin Resistant Staphylococcus aureus. Front Microbiol. 2020;11:695.

