## Supplemental for "Combining Bacteriophage and Vancomycin is Efficacious Against MRSA biofilm-like Aggregates Formed in Synovial Fluid"

**Supporting Information**

**Table S1.** *Galleria mellonella* health index scoring system.

| Category | Observation | Score |
|---|---|---|
| Activity | No activity | 0 |
| | Minimal response to stimuli | 1 |
| | Active response to stimuli | 2 |
| | Active without stimulation | 3 |

**Table S2.** InterProScan functional domain results for phage Remus

| Gene product; protein ID | Name | Type | Minimum (AA) | Maximum (AA) | Length (AA) |
|---|---|---|---|---|---|
| 134; YP_008431175.1 | Coil | COILS | 96 | 116 | 21 |
| | DUF2977 | Pfam | 14 | 74 | 61 |
| | hypothetical protein CDS | CDS | 1 | 119 | 119 |
| 137; YP_008431172.1 | DUF4815 | Pfam | 9 | 611 | 603 |
| | DUF4815 domain-containing protein CDS | CDS | 1 | 1153 | 1153 |
| 138; YP_008431171.1 | structural protein CDS | CDS | 1 | 174 | 174 |
| | baseplate structural protein gp8, domain 1 | Gene3D | 6 | 106 | 101 |
| 140; YP_008431169.1 | baseplate J/gp47 family protein CDS | CDS | 1 | 349 | 349 |
| | Baseplate_J | Pfam | 66 | 308 | 243 |
| 141; YP_008431168.1 | baseplate protein CDS | CDS | 1 | 235 | 235 |
| | LysM | CDD | 6 | 41 | 36 |
| | LysM_2 | SMART | 5 | 66 | 62 |
| | gpW/gp25-like | Superfamily | 125 | 212 | 88 |
| | | Gene3D | 110 | 221 | 112 |
| | LysM domain | Gene3D | 3 | 76 | 74 |
| 144; YP_008431165.1 | glycerophosphoryl diester phosphodiesterase CDS | CDS | 1 | 819 | 819 |
| | Phosphatidylinositol (PI) phosphodiesterase | Gene3D | 601 | 818 | 218 |
| | GP_PDE | PROSITE_PROFILES | 593 | 818 | 226 |
| | GLYCEROPHOSPHORYL DIESTER PHOSPHODIESTERASE | Panther | 603 | 817 | 215 |
| | GDPD | Pfam | 606 | 812 | 207 |
| | PLC-like phosphodiesterases | Superfamily | 618 | 816 | 199 |
| | Coil | COILS | 406 | 426 | 21 |
| | Coil | COILS | 349 | 369 | 21 |
| | Coil | COILS | 316 | 336 | 21 |

| | | | | | |
|---|---|---|---|---|---|
| 145; YP_008431165.1 | endopeptidase domain like (3.90.1720.10) | Gene3D | 151 | 293 | 143 |
| | NLPC_P60 | PROSITE_PROFILES | 146 | 290 | 145 |
| | NLPC_P60 | Pfam | 164 | 259 | 96 |
| | Cysteine proteinases | Superfamily | 156 | 289 | 134 |
| | tail lysin CDS | CDS | 1 | 297 | 297 |
| 146; YP_008431163.1 | endopeptidase domain like (3.90.1720.10) | Gene3D | 688 | 807 | 120 |
| | CHAP | PROSITE_PROFILES | 676 | 807 | 132 |
| | CHAP | Pfam | 695 | 781 | 87 |
| | Cysteine proteinases | Superfamily | 687 | 795 | 109 |
| | disorder_prediction | MOBIDB_LITE | 645 | 670 | 26 |
| | disorder_prediction | MOBIDB_LITE | 645 | 662 | 18 |
| | CHAP domain-containing protein CDS | CDS | 1 | 811 | 811 |
| 147; YP_008431162.1 | hypothetical protein CDS | CDS | 1 | 277 | 277 |
| | Coil | COILS | 18 | 38 | 21 |
| | 1.10.530.10 | Gene3D | 52 | 196 | 145 |
| | Phage_lysozyme2 | Pfam | 61 | 196 | 136 |
| | disorder_prediction | MOBIDB_LITE | 35 | 59 | 25 |
| | disorder_prediction | MOBIDB_LITE | 1 | 20 | 20 |
| | disorder_prediction | MOBIDB_LITE | 1 | 15 | 15 |
| 148; YP_008431161.1 | 1.10.530.10 | Gene3D | 115 | 231 | 117 |
| | Glucosaminidase | Pfam | 120 | 229 | 110 |
| | Lysozyme_4 | SMART | 106 | 242 | 137 |
| | disorder_prediction | MOBIDB_LITE | 18 | 40 | 23 |
| | disorder_prediction | MOBIDB_LITE | 18 | 37 | 20 |
| | tail tip protein CDS | CDS | 1 | 290 | 290 |

**Figure S1**

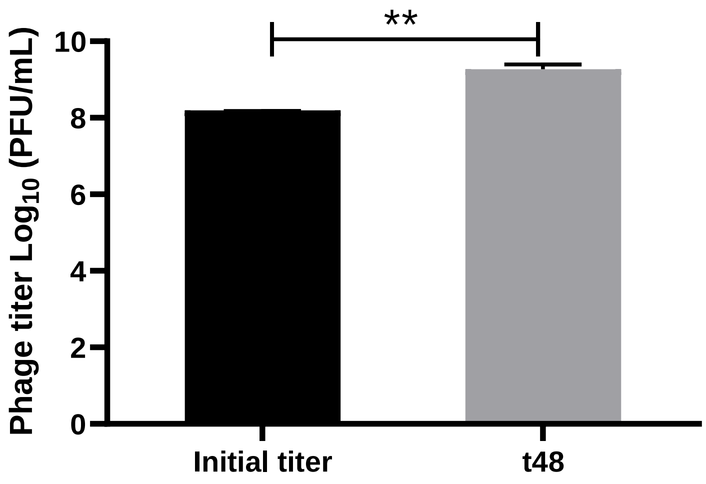


**Figure S2**

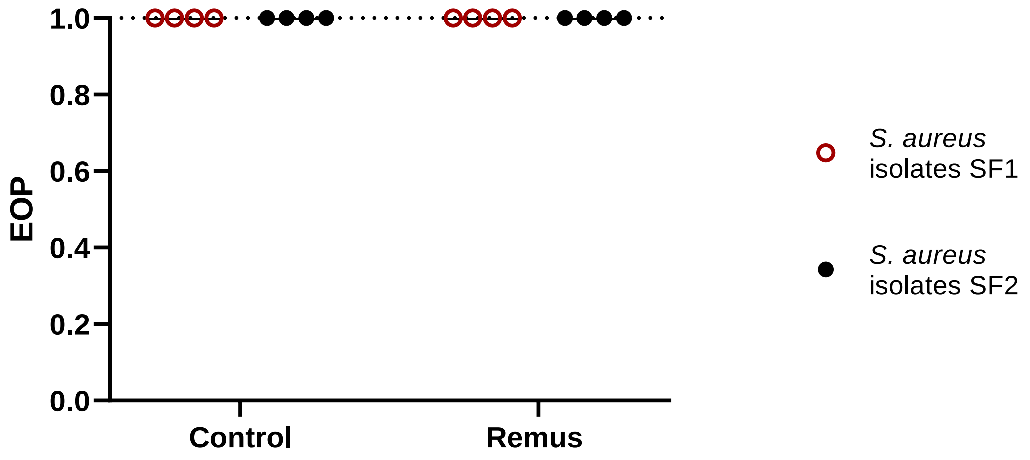
